# Concomitant phytonutrient and transcriptome analysis of mature fruit and leaf tissues of tomato (*Solanum lycopersicum* L. cv. Oregon Spring) grown using organic and conventional fertilizer

**DOI:** 10.1101/755769

**Authors:** Richard M Sharpe, Luke Gustafson, Seanna Hewitt, Benjamin Kilian, James Crabb, Christopher Hendrickson, Derick Jiwan, Preston Andrews, Amit Dhingra

**Author notes:** **Corresponding Author:** Amit Dhingra, Ph.D., Department of Horticulture, Washington State University, PO Box 646414, Pullman, WA 99164-6414.

## Abstract

Enhanced levels of antioxidants, phenolic compounds, carotenoids and vitamin C have been reported for several crops grown under organic fertilizer, albeit with yield penalties. As organic agricultural practices continue to grow and find favor it is critical to gain an understanding of the molecular underpinnings of the factors that limit the yields in organically farmed crops. Concomitant phytochemical and transcriptomic analysis was performed on mature fruit and leaf tissues derived from *Solanum lycopersicum* L. ‘Oregon Spring’ grown under organic and conventional fertilizer conditions to evaluate the following hypotheses. 1. Organic soil fertilizer management results in greater allocation of photosynthetically derived resources to the synthesis of secondary metabolites than to plant growth, and 2. Genes involved in changes in the accumulation of phytonutrients under organic fertilizer regime will exhibit differential expression, and that the growth under different fertilizer treatments will elicit a differential response from the tomato genome. Both these hypotheses were supported, suggesting an adjustment of the metabolic and genomic activity of the plant in response to different fertilizers. Organic fertilizer treatment showed an activation of photoinhibitory processes through differential activation of nitrogen transport and assimilation genes resulting in higher accumulation of phytonutrients. This information can be used to identify alleles for breeding crops that allow for efficient utilization of organic inputs.

**Significance statement:** Organic fertilizer changes the expression of the tomato genome, induces photosynthetic stress which elicits higher production of secondary metabolites.

## INTRODUCTION

As agriculture prepares to support 10 billion people by 2050, grow 70% more food in increasingly less amount of land and reduce environmental impact [1], there is a need to assess different modes of farming. The proponents of organic farming have stated that it can have reduced environmental impact from savings in fossil fuels, reduced environmental pollution, and fostering of greater biodiversity however, these benefits accrue at a cost of reduced yield which can range from 5-34% depending on the conditions used [2–4].

A recent meta-analyses, based on data from 343 peer-reviewed publications using a standardized weighted analysis, concluded that on an average, organically grown crops have higher concentrations of antioxidants and a lower percentage of pesticide residues compared to conventional crops across regions and production seasons [5]. This report is in contrast to some previously reported analyses that found no significant difference between the foods grown in conventional vs. organic systems [6, 7]. Several studies have reported the biochemical analyses of organically grown apple, strawberry, and tomato fruits where the levels of antioxidants, phenolic compounds, carotenoids and vitamin C were seen to be enhanced. [8–10]. Based on the incidence of higher phytonutrient content, organically produced foods have also been proposed to be more nutritious compared to their conventionally grown counterparts however, the scientific opinion remains divided [11].

Organic farming necessitates that the nitrogen needs of the plant are met from manures derived from animal byproducts, crop residues, green manures, legumes and soil organic matter [12]. However, the process of developing new crops via plant breeding utilizes conventional agricultural practices, thereby selecting crops with an inherent genetic bias towards utilization of conventional fertilizers and management practices [13]. The fundamental difference between conventionally or organically grown foods is the chemical nature of the nutrition inputs that a plant must utilize. The presence of carbon-linked inorganic nutrients impacts the bioavailability of the inorganic elements. It is assumed that microbial communities first degrade the organic nutrients, which are then absorbed by the plant [14]. Therefore, organic production systems are far more complex than conventional systems and the nutrition of the plant must depend more on the available microbial communities. This begs the following questions: Do the plants, having been bred under conventional inputs, adjust their response at the genetic level to utilize organic nutritional inputs, and interactions with available microbial communities? Is there a natural degree of malleability rendered by some genetic backgrounds to adapt better to the organic inputs? Are there plant metabolic pathways that could be altered with external, but organic, inputs to overcome the yield losses reported under organic conditions?

A very limited number of gene expression studies have attempted to address the aforementioned questions to gain valuable insights into plant’s genomic and metabolic performance under organic and conventional fertilizer regimens. A microarray based gene expression study in potato was used to identify statistically different expression profiles under organic and conventional treatments [15]. Further, based on different gene ontology pathways, differentially expressed genes were also identified [16]. In wheat, univariate and multivariate statistical analysis was done on microarray-based gene expression data to establish that the organic inputs influence the expression of global wheat transcriptome, which could be utilized to verify the production system at the farm level [17]. Perhaps, due to the confounding impact of the environment, none of these reports were able to isolate and quantify the impact of organic inputs on a plant’s metabolic and transcriptomic response. Furthermore, none of the previous studies reported changes in both the phytochemical composition and the concomitant changes in the global gene expression.

The major concern with organic inputs is the losses in yield which are counter to what is needed to feed a burgeoning population [18]. However, the organic agricultural practices continue to grow and find favor with customers making it both critical and urgent to gain an understanding of the molecular underpinnings of the factors that limit the yield in organically farmed crops. It has been proposed that with proper management practices, crops yields obtained under organic farming can match conventionally farmed crops [19]. Understanding the molecular basis can help in identifying proactive and predictive agronomic, agrochemical or biological interventions that can enhance crop productivity in organic systems. This can have significant implications in enabling small-holding farms across the globe, which have relatively easier access to organic inputs compared to conventional inputs, to grow food sustainably.

In order to understand the impact of conventional and organic inputs on a single genotype of tomato, without the confounding factor of the environment, this study was conducted under controlled environmental conditions. Two hypotheses were evaluated in this study: 1. Organic soil fertilizer management, which results in a slower rate of biological release of available nitrogen to plant roots, results in greater allocation of photosynthetically derived resources to the synthesis of secondary metabolites, such as phenolics and other antioxidants, than to plant growth, and 2. Genes involved in changes in the accumulation of phytonutrients under organic fertilizer regime will exhibit differential expression, and that the growth under different fertilizer treatments will elicit a differential response from the tomato genome. Both these hypotheses were supported, suggesting an adjustment of the plants’ metabolic and genomic activity in response to different nitrogen regimes.

## RESULTS AND DISCUSSION

### Agronomic and Phytonutrient analysis

The mean mass of red ripe fruit was 20% greater in conventionally (CONV) grown plants (Supplementary Figure 1). Similarly, the cumulative above and below ground vegetative biomass on a fresh weight basis was approximately 4% greater in CONV grown plants. While the above ground vegetative biomass on a dry weight basis was 4.5% higher in CONV grown plants, the below ground vegetative biomass on a dry weight was 6% greater in organically (ORG) grown plants (Supplementary Table 1). The observed reduction in yield and biomass under ORG fertilizer is as expected since the use of organic fertilizer has been implicated in the reduction of yield by 5-34% [4].

Total soluble solids were higher by 18.9% in ripe ORG fruit, which also had higher concentrations of phenolic compounds (FW – 17.02%, DW – 16.81%), lycopene (FW and DW – 11.93%), and vitamin C (FW – 11.46%, DW – 13.88%) compared to CONV fruits (**Table 1**). Leaf C: N ratio was higher by 12.5% in ORG plants, thus most likely favoring the synthesis of C-based compounds, like phenolics, ascorbic acid, and carotenoids (Supplementary Table 2). Similar results regarding accumulation of phytonutrients has been reported in several previous studies [9, 10]. The observation of enhanced levels of phytonutrients supported the first hypothesis that growth of tomato plants under organic fertilizer management results in greater accumulation of secondary metabolites, such as phenolics and other antioxidants, than to plant growth. This may be due to the preferential allocation of photosynthetically derived resources to the synthesis of these secondary metabolites.

**Table 1:**
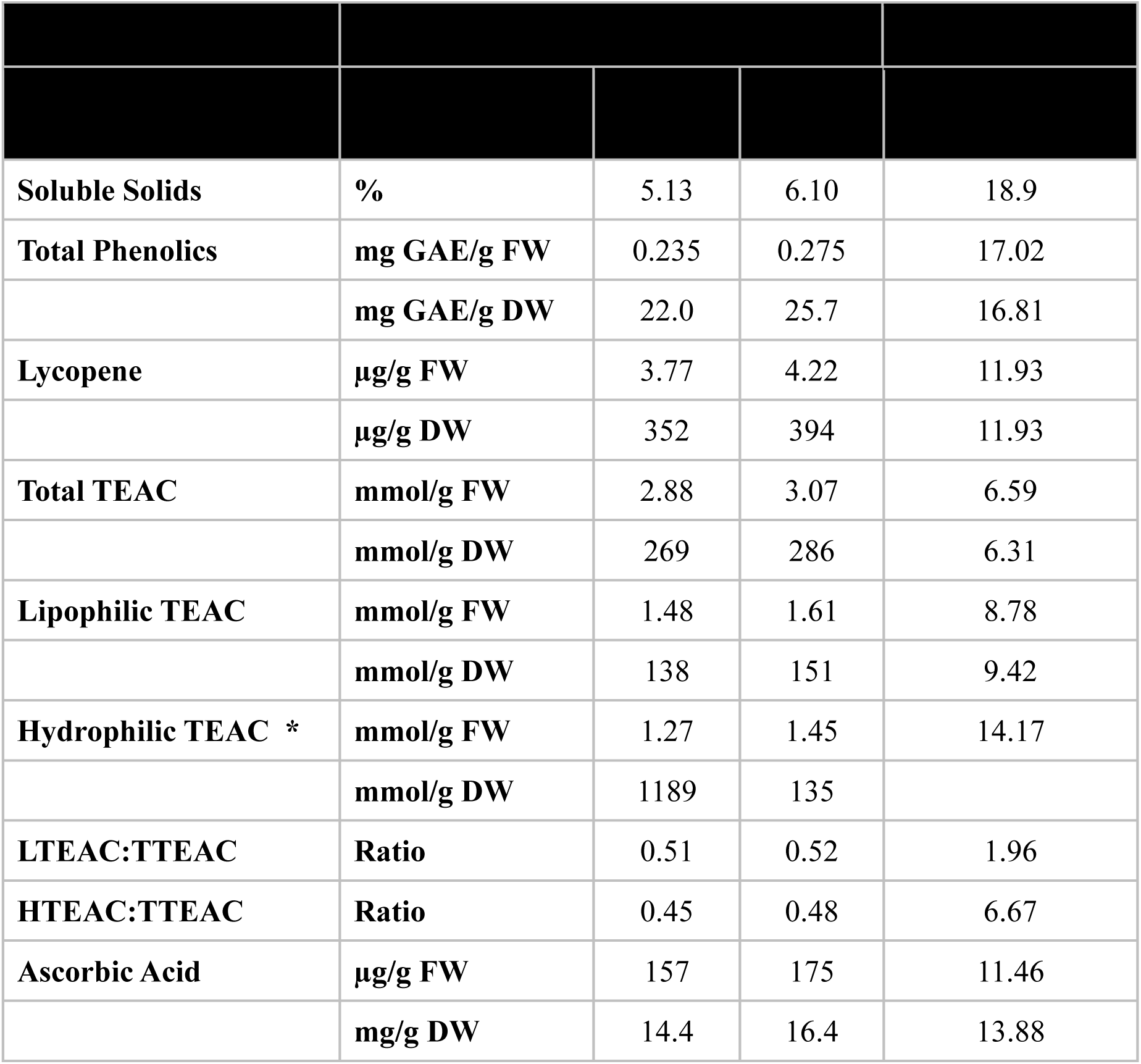
Measured phytonutrient content. Soluble solids, total phenolics (gallic acid equivalents, GAE), lycopene, total Trolox equivalent antioxidant capacity (TTEAC), lypophilic TEAC, hydrophilic TEAC, and reduced ascorbic acid concentrations on fresh (FW) and dry weight (DW) bases (except soluble solids) under conventional (CONV) and organic (ORG) fertility treatments. Data were analyzed using ANOVA mixed model. Failure to meet normality based on Kolmogorov-Smirnov test is indicated by asterisk (*).

### RNAseq and biochemical pathway analysis

#### Transcriptome summary

A tabulation of total raw sequence reads, number of curated reads, number of assembled contiguous sequences (contigs) or expressed sequence tags (ESTs), and number of ESTs exhibiting very high differential expression ratios, equal to or greater than 5-fold expression difference, between the ORG and CONV treatments for each tissue and treatment is summarized in Supplementary Table 3. A total of 2,324 and 3,035 contigs were identified to exhibit equal to or greater than 5-fold expression difference between the ORG and CONV treatments in fruit and leaf samples, respectively.

#### Quantitative RT-PCR verification

The expression trends obtained from the RPKM values were verified by performing qRT-PCR based expression of 27 selected genes. Of these, eight genes were selected to be evaluated as reference genes based upon the fact that their tissue and sample specific RPKM ratios were nearly zero, which is indicative of equivalent expression between the tissue and sample types. From these eight candidate genes, Calreticulin3-like, accession number Solyc05g056230.3.1, was selected as the reference gene due to the equivalent tissue specific expression values obtained from RPKM and qRT-PCR results (Supplementary Figure 2).

#### Functional annotation

The transcriptome data was processed using Blast2GO sequence alignment, gene ontology (GO) mapping, and functional annotation workflow [20–22]. Sequences were processed through BLAST against the Viridiplantae database using an e-value cutoff of 1.0e−3. After Blast2GO processing, the relative expression of contigs (RPKM values) with GO terms involved in the differential accumulation of phytonutrients was analyzed.

### Global changes in gene expression under CONV and ORG conditions

Observed differences in the phytonutrient and agronomic characteristics of tomato plants grown under ORG and CONV fertility treatments (Section 1) indicated that the growth under different fertilizer treatments are expected to elicit a differential expression response at a global level. Visualization of the log10 ratio of RPKM values (ORG/CONV) across each chromosome of tomato reflected the difference in expression. The observed shift of the trend lines, indicative of a predominance of overall chromosomal expression either in the ORG treatment (positive y-intercept value) or in the CONV treatment (negative y-intercept value), indicated there was an enhanced expression at certain genomic loci in ORG grown plants (**Figure 1** and Supplementary Figures 3A and 3B). This observation supports a part of the second hypothesis that growth of the same genotype of tomato under different fertilizer treatments will elicit a differential expression response from the tomato genome.

**Figure 1.**
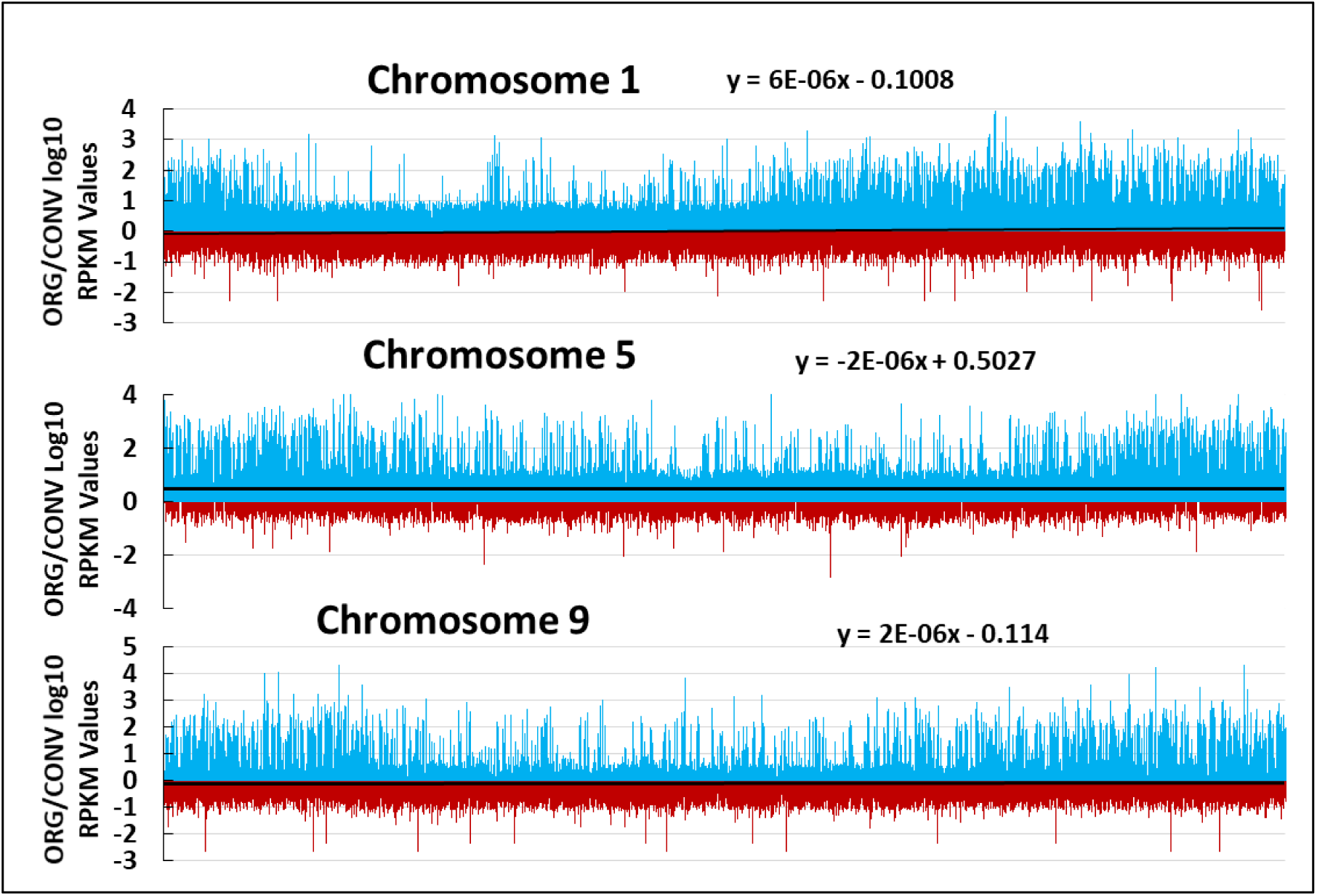
Representative ORG and CONV log10 RPKM ratio Manhattan plot representations of expression values across *S. lycopersicum* chromosomes. Every bar is indicative of the log10 expression ratio of Fruit and Leaf ORG RPKM values over Fruit and Leaf CONV RPKM values. Trendlines with slope equations indicate propensity of treatment activated loci across the chromosome. Chromosome graph lengths are not indicative of chromosome length. Red bars indicate a higher log10 ratio in the CONV treatment and blue bars indicate higher expression in the ORG treatment at any given position on the chromosome. Trend lines were graphed to indicate a predominance of chromosomal activity either in the ORG treatment (positive y-intercept value) or in the CONV treatment (negative y-intercept value). All 12 mapped chromosomes can be found in Supplemental Figures 3a and 3b. 12 read files, representing three cDNA libraries across each of the two tissue and two treatment types were used for mapping purposes.

### Gene Ontology and KEGG Pathway analysis related to Phytonutrient biosynthesis and Primary Metabolic pathways

Several recent studies have utilized transcriptomics-based gene ontology and Kyoto Encyclopedia of Genes and Genomes (KEGG) pathway analysis for establishing a theoretical basis to understand the metabolic framework underlying complex developmental processes or organismal responses to external cues in tomato and other species [23–28].

In this study, GO and KEGG pathway analysis of the transcriptome data focused on genes related to biosynthesis of phytonutrients that were observed to accumulate differentially under the two conditions. Prior studies have attributed the observed changes in metabolites primarily to stress, however, the role of nitrogen source was left unaddressed [29, 30].

Besides the organic forms of nitrogen, organic fertilizer predominantly contains amino acids and ammonia, which is known to be toxic to tomato [31]. Efforts to overcome the toxicity of ammonia likely activates the salvage pathways that include photorespiratory processes and production of phytonutrients protective to the plant. Since ammonia impacts nitrogen metabolism and photosynthetic processes in the plant [32], the biochemical pathway analysis of the transcriptome data also included an evaluation of the two primary metabolic pathways to gain an understanding of their impact and interaction when the plants were grown under the two fertilizer conditions.

### Impact of ORG inputs on Lycopene and Ascorbate pathways

#### Lycopene

An approximate increase of 12% in Lycopene content was observed in the fruit, both on a fresh and dry weight basis under ORG fertilizer (Table 1). Analysis of the expression values of all the genes involved in the Lycopene pathway revealed that Phytoene synthase 1 Solyc03g031860.3.1, representing the first committed enzyme in the lycopene pathway, was the highest expressed transcript in the pathway in the fruit tissue compared to the leaf tissue, however there were no differences in the fruit transcripts from the two treatments. Interestingly, the expression of Phytoene synthase 1 (*Psy1*) was higher in the ORG leaf tissue compared to the CONV leaf tissue (**Figure 2**). Phytoene synthase 2 (*Psy2*) (Solyc02g081330.4.1), a chloroplast targeted specific homolog exhibited an overall low expression in both tissues, and did not indicate a difference between the treatment types (Supplementary Figure 4).

**Figure 2.**
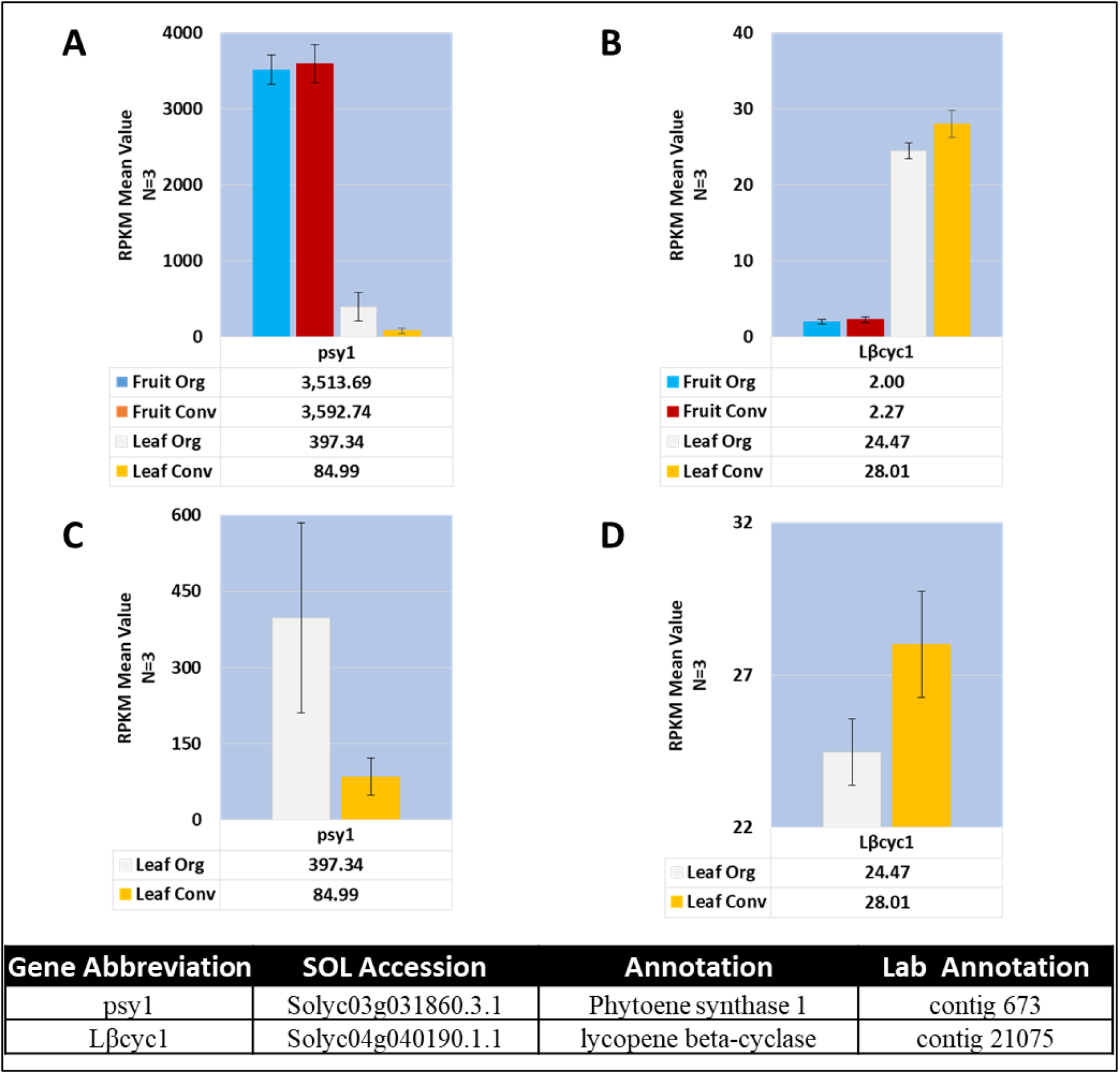
Mean expression profile of phytoene synthase 1 (*psy1*) and lycopene beta-cyclase (*Lβcyc1*). PSY1 is the rate limiting step in the carotenoid biosynthesis pathway and the absence of LβCYC in the pathway increases the pool of lycopene from the pathway. **A.** *psy1* RPKM expression values. **B.** *Lβcyc1* RPKM expression values. **C.** *psy1* leaf expression values with a different y-axis to discern standard error differences. **D.** *Lβcyc1* leaf expression values with a different y-axis to discern standard error differences. Table of annotations and accession numbers from Sol Genomics ITAG3.2. Error bars denote the standard error of sequenced expression values between 3 biological replicates.

The first question that emerges from these observations is what is contributing to the higher levels of lycopene in the fruit? Capture of transcriptome expression at the mature stage does not provide any relevant information. Perhaps, the higher accumulation could be attributed to higher activity of Psy1 or other enzymes involved in the pathways during breaker or earlier stages of fruit development.

The second question is why the levels of *Psy1* are higher in ORG leaves and what function may they be serving? Psy1 is active primarily in chromoplasts, and Psy2 is a chloroplast predominant enzyme [33]. The lycopene pool fluxes through lycopene β-cyclase, in the lycopene B pathway producing β-carotene, antheraxanthin, violaxanthin and neoxanthin, or through lycopene ε-cyclase in the lycopene A pathway producing α-carotene and lutein [33]. Upregulation of *Psy1* in the leaf under ORG conditions is intriguing. One explanation could be that the additional activity of *Psy1* is resulting in the production of additional pools of lycopene that may be converted into carotenoid or ABA via activation of NCED converting violaxanthin to xanthoxin to compensate for photorespiratory stress [33].

Lycopene cyclases, enzymes responsible for converting lycopene metabolite pools for downstream carotenoid production, were down regulated in the fruit tissue compared to the leaf tissue, which, as expected is responsible for higher lycopene content in the fruit tissue compared to leaf (Figure 2 and Supplementary Figure 4). Increased lycopene cyclase activity, as well as between the beta and epsilon isoforms, in tomato fruit has been shown to decrease lycopene content [34, 35]. Absence of differential activity of *Psy1* and lycopene cyclases in the ORG and CONV fruit tissues indicates that there is a need for a developmental time course transcriptome analysis under these two conditions to identify the molecular reason underlying higher observed lycopene content in ORG fruit.

#### Biosynthesis and recycling of ascorbate

In tomato the major pathway leading to accumulation of Ascorbic acid (AsA) is the Smirnoff-Wheeler pathway [36], whereas the recycling of AsA to control oxidative stress is regulated by the Foyer-Halliwell-Asada pathway [37]. Transcripts representing the enzymes involved in biosynthesis and recycling of AsA were assessed for their expression levels.

#### Biosynthesis of AsA - Smirnoff-Wheeler pathway

Depending on the plant species, transcriptional activity of two genes has been reported to be directly implicated in the overall production of AsA [38]. In Arabidopsis and kiwi fruit, GDP-L-galactose phosphorylase/guanyltransferase (GGP) was identified as a rate limiting step for synthesis of AsA [39]. In ripening tomatoes, the overall production of AsA was reported to be dependent on the expression of L-galactose-1-phosphate phosphatase (GPP) [40]. However, overexpression of GGP resulted in a 3- to 6-fold increase of AsA in transgenic tomato fruit indicating GGP expression can also contribute to total AsA in tomato fruit [41].

In this study, less than a 9% difference in the transcriptional abundance of *GPP1* between the fruit tissues was observed (**Figure 3**, Supplementary Figure 5 and Supplementary Table 4). However, one of the isoforms of *GGP* had the highest expression in the CONV leaf tissue while *GGP* transcript abundance was 25% less in the ORG leaf (Supplementary Figure 5). Similarly, two genes involved in Ascorbate synthesis, phosphomannose mutase (PMM) and L-galactose dehydrogenase (galDH), for which a single isoform was identified, expressed at similar levels in both tissue types and treatments (Supplementary Figure 5). The expression of genes representing phosphoglucose isomerase (*PGI*) and L-galactono 1,4-lactone dehydrogenase (*galLDH*), were upregulated in a tissue specific manner. Two isoforms of *PGI* were identified and they both expressed at a higher level in the fruit but expressed at a similar level in both the treatments. There was a single isoform of *galLDH* that was upregulated in the leaf tissue but expressed within one RPKM value between treatments (Supplementary Figure 5).

**Figure 3.**
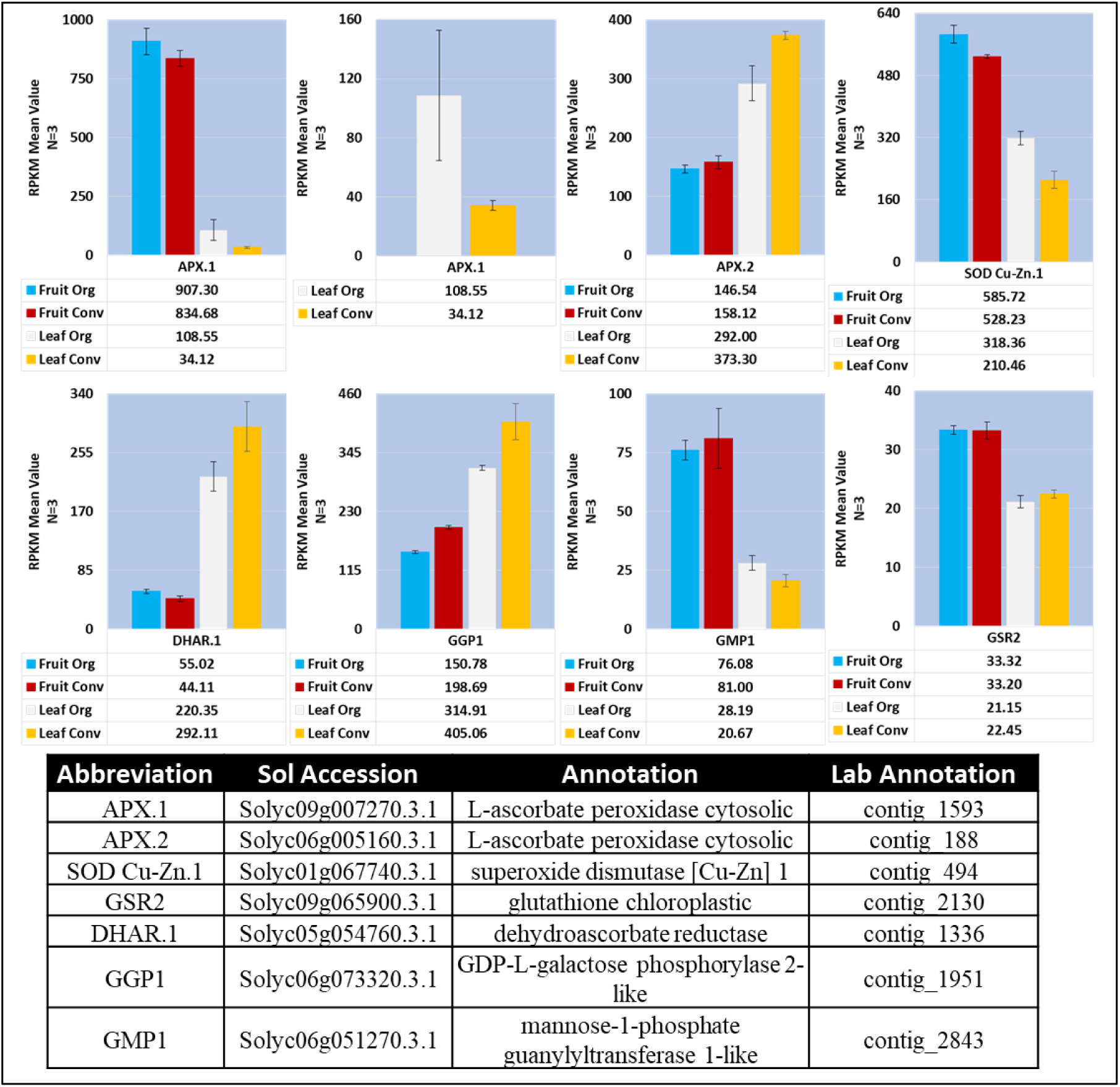
Mean expression profile of significant ascorbate pathway enzymes. GGP1 and GMP1 are involved in the production of ascorbate (Smirnoff-Wheeler Pathway). APX.1, APX.2, SOD Cu-Zn.1, GSR2 and DHAR.1 are involved in the activation of ascorbate (Foyer-Halliwell-Asada Pathway). Table of annotations and accession numbers from Sol Genomics ITAG3.2. Additional ascorbate pathway gene expression can be found in supplementary figures 5, 6a and 6b. Error bars denote the standard error of sequenced expression values between 3 biological replicates.

Three GDP-D-mannose phosphorylase (*GMP*) isoforms were identified in the transcriptome where the highest RPKM values were reported in the fruit tissue for one of the *GMP* isoforms. The other two isoform RPKM values were approximately 75% of the previously described highest fruit isoform. One of the *GMP* isoforms was equally expressed between the tissue and treatment types and the third isoform was expressed at a higher level in the leaf tissue. None of the three isoforms were differentially expressed between the ORG and CONV treatments (Supplementary Figure 5).

GDP-D-mannose 3,5-epimerase (*GME*) expression indicated the presence of two isoforms with one isoform expressed higher in the fruit tissue by approximately 66% and the second isoform expression trend towards a higher leaf expression value. Neither of the two isoforms were differentially expressed between the ORG and CONV treatments. The final gene identified in the Smirnoff-Wheeler pathway, mannose-6-phosphate isomerase (*MPI*) was represented by two different isoforms where neither isoform was differentially expressed in the ORG and CONV treatments. One isoform did not indicate differential expression values between the tissue types while the other isoform RPKM values were higher in the leaf tissue (Supplementary Figure 5).

#### Recycling of AsA - Foyer-Halliwell-Asada pathway

The metabolic functions of ascorbate in plant tissue require a recycling pathway referred to as the Foyer-Halliwell-Asada pathway [42]. The expression of genes coding for the Foyer-Halliwell-Asada enzymes was evaluated (**Figure 3**, Supplementary Figure 6 and Supplementary Table 5). Ten of the 21 identified genes, gene isoforms or organelle targeted specific isoforms in the pathway were expressed at a higher level in a tissue specific manner (**Figure 3** and Supplementary Figure 6). In the transcriptome dataset, one each of a cytosolic and a mitochondrial isoform, and two peroxisomal monodehydroascorbate reductase (*MDHAR*) isoforms were identified. All *MDHAR* isoforms were expressed at a higher level in the fruit tissue. Except for the cytosolic isoform (Solyc09g009390.3.1) that was expressed at approximately 65 percent of the leaf levels, other isoforms were expressed at approximately 2-fold higher level in the fruit tissue. However, none of the isoforms demonstrated differential expression between the ORG and CONV treatments in any of the tissue types.

As a part of the reduction/oxidation of glutathione branch of the pathway [42], two isoforms of glutathione reductase (*GSR*) (Solyc09g091840.3.1 and Solyc09g065900.3.1) and one isoform of dehydroascorbate reductase (*DHAR*) (Solyc05g054760.3.1) were identified. In the leaf tissue, an isoform each of *GSR* and *DHAR* demonstrated higher expression values in the CONV treatment in comparison to the ORG treatment although the difference was less than two-fold (**Figure 3**). Another isoform of *GSR* was expressed at a lower level with no discernable differences between the ORG and CONV treatments.

Interestingly, evaluation of the expression levels of the enzymes responsible for the cycling of redox status of glutathione in the Foyer-Halliwell-Asada cycle indicate that the greatest transcript activity occurs in the leaf tissue. Enzymes responsible for the detoxification of H2O2 - members of the ascorbate peroxidase and superoxide dismutase families - were expressed at a higher level in the fruit tissue, except for one cytosolic ascorbate peroxidase (*APX*)(Solyc06g005160.3.1) transcript, and a peroxisomal APX (*peroxAPX*)(Solyc01g111510.3.1). A cytosolic *APX* transcript (Solyc09g007270.3.1) demonstrated a 10-fold higher expression in fruit than the leaf. The expression was higher in both the leaf and fruit tissues from ORG treatment compared to CONV treatment (**Figure 3**). Four additional *APX* isoforms, two *APX6* and two APX6 variant 2 (*APX6 x2*), were also identified as having a relatively low expression with one *APX6* and one *APX6 x2* isoforms expressed at a higher level in the fruit tissue (Supplementary Figure 6).

A cytosolic superoxide dismutase (*SOD*) transcript belonging to the copper-zinc (Cu-Zn) family (Solyc01g067740.3.1) demonstrated 2-fold higher expression values in the fruit compared to the leaf. The expression was higher in the ORG treatment (**Figure 3**). Additional *SOD* transcripts were also detected; one *SOD Cu-Zn*, a chloroplast targeted SOD Cu-Zn (*cpSOD Cu-ZN*), four chloroplast targeted SOD Fe family isoforms and a mitochondria targeted SOD Mn family isoform (Supplementary Figure 6 and Supplementary Table 5). All the additional *SOD* isoforms were expressed at a higher level in the fruit tissue or equally between the fruit and leaf tissues with the exception of a *cpSOD Fe* isoform. Notably, there was one *cpSOD Cu-Zn* transcript whose expression values were higher in the fruit tissue. While the expression was comparable between the ORG and CONV treatments, the leaf ORG expression values were almost four times compared to the leaf CONV treatment (Supplementary Figure 6B).

An obvious correlation between the observed levels of ascorbate in the fruit tissue with the expression patterns of genes coding for enzymes in the ascorbate biosynthesis and recycling of ascorbate is missing. This is most likely due to the fact that in this study mature fruit tissues were used for the transcriptome analysis necessitating a time course developmental biochemical and transcriptome analysis of the fruit tissue to understand the role of all the genes involved in ascorbate biosynthesis. Higher overall expression of transcripts coding for enzymes involved in recycling in the ORG leaf tissues is interesting. It indicates that ascorbate recycling may be operating at a higher level under ORG conditions and may contribute to higher ascorbate levels in the fruit. Although fruit microclimate has been shown to influence fruit ascorbate levels as well [43, 44].

#### Impact of ORG inputs on biosynthesis of TEAC, total phenolics, and soluble solids

Three of the five phytonutrients quantified in this study, namely TEAC, total phenolics, and soluble solids, represent a combination of metabolic components rendering it difficult to ascribe the contribution of specific enzymes towards the observed final concentration. For instance, the TEAC test for antioxidant capacity does not specifically test for individual antioxidants, but rather total antioxidant capacity, and should not be used to identify individual antioxidants by kinetics or stoichiometry [45]. As discrete antioxidant activity cannot be ascertained with this method, it was not feasible to analyze the expression of genes coding of enzymes involved in these pathways. Similar to the TEAC test, the Folin-Ciocalteu assay for total phenolic content assessment [46], reacts with at least 33 different phenols or compounds when the assay is performed on total tissue eluates [46, 47]. Therefore, the assay could not be used to quantify levels of any particular phenol without specific metabolite assays. Sugars, organic acids, amino acids and soluble pectins constitute the majority of soluble solids reported as °Brix values from refractometer measurements. Given the complexity of the metabolites being detected for the three classes of phytonutrients, it would not be informative to draw any correlations to specific genes that participate in the biosynthetic pathways of these phytonutrients. Therefore, Fisher’s exact test was utilized as an alternative approach to identify the differential expression of genes involved in these pathways.

Comparison between the combined Leaf ORG and CONV treatments (leaf subset) and the combined reference yielded no significant GO term enrichment. Since there was no enrichment of GO terms in the leaf tissue, it should not be assumed that the response of individual transcripts in the leaves were similar under the two conditions. This is illustrated by the RPKM values of *Psy1*, which were 3.2-fold higher in the ORG treatment versus the CONV treatment (**Figure 2**).

However, interestingly, among the combined fruit ORG and CONV (fruit subset) comparison, terms associated with the chloroplast, photosynthesis, organic acid cycling processes and functions of the plastids, specifically the chloroplast, thylakoid and photosystems, and sulfur compound biosynthesis were over represented in the Fruit ORG tissues (**Figure 4**). In the Fruit ORG treatment there was a higher representation of genes related to photosynthesis and photosynthetic stress. The underrepresented fruit ORG GO terms included non-specific organelle, structural molecule activity and translational machinery (**Figure 4**). The underrepresented GO terms in the ORG fruit were involved in the cellular component ontology, mainly involved in cell structure components. Underrepresented GO terms in the three ontology categories represented greater than half of the contigs that showed a lower expression of 2-fold or lower values with the exception of three GO terms.

**Figure 4.**
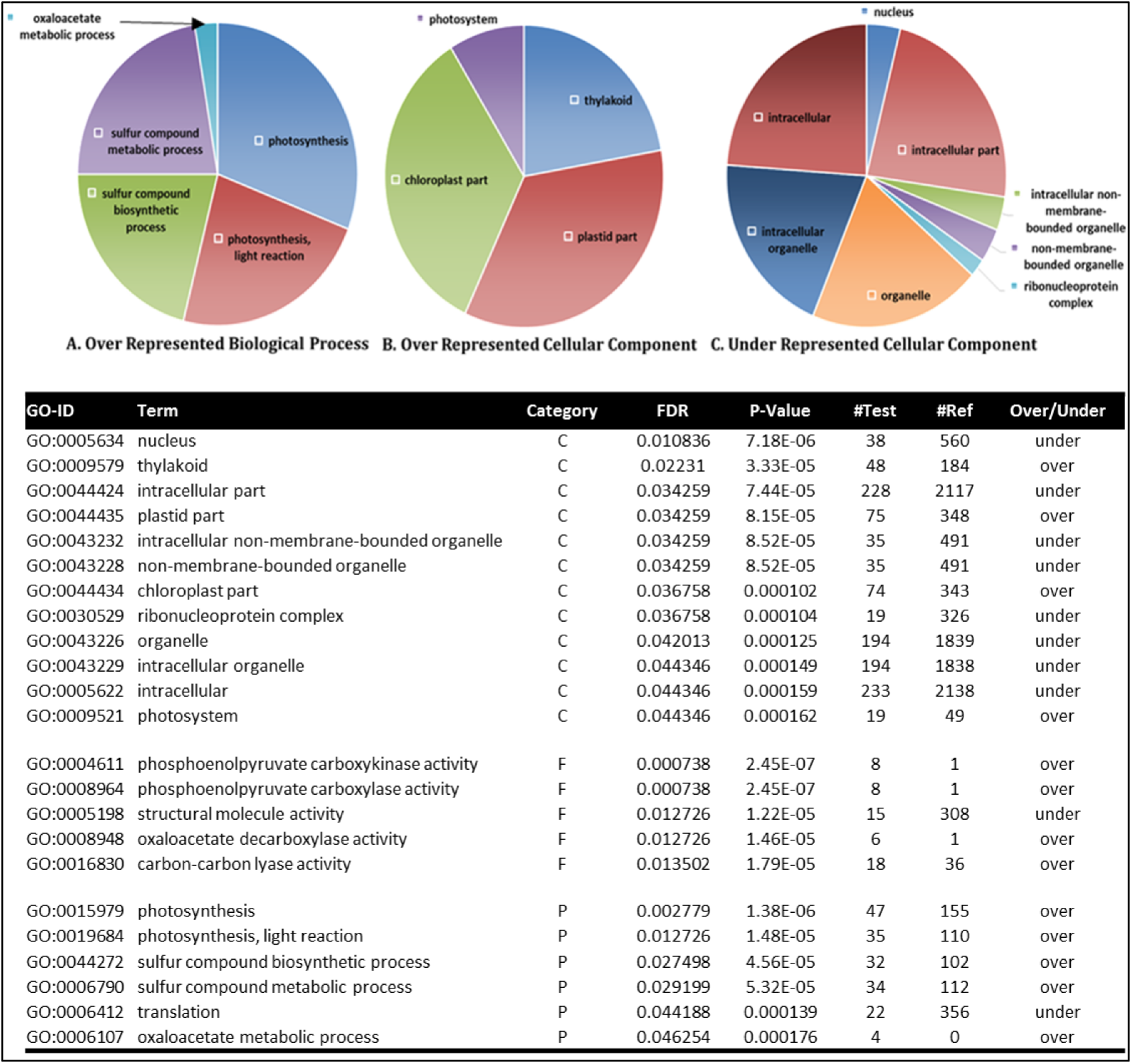
Gene Ontology Fisher’s Blast2GO analysis and queried with fruit tissues. **A.** Biological Process gene ontology terms over represented in the ORG treatment compared to CONV treatment. **B**. Over represented Cellular Components in the ORG treatment versus the CONV treatment. **C.** Cellular Component ontology terms under represented in the ORG treatment compared to the CONV treatment. Size of slices represent ratio of numbers of contigs to total number of contigs in the represented categories.

When the contigs corresponding to the enriched GO terms were mined and their RPKM values compared, it was revealed that all of them demonstrated a 2-fold greater RPKM value for the ORG treatment than either the CONV or non-differential RPKM values (Supplementary Figure 7). One exception was GO: 0043226 organelle cellular component, which had a larger proportion of contigs that were non-differentially expressed (Supplementary Figure 7). While the number of contigs expressed at 2-fold or higher level in the ORG treatment for two of the terms, GO:0005622 and GO:0005634 comprised 50% of the contigs, one term, GO:0043226 was comprised of more non-differentially expressed contigs with 34.6% of the contigs expressed at 2-fold or higher level in the ORG treatment (Supplementary Figure 7).

GO term enrichment between the differentially expressed leaf and fruit ORG subset and the leaf and fruit CONV subset also returned greater ratio of contigs represented in the ORG subset. Surprisingly, 10 of the 23 ontology terms with higher ORG ratios included under-represented GO terms, i.e. GO:0005622, GO:006412 and GO:0005198, and the Cellular Component ontology had two times more underrepresented GO terms than overrepresented GO terms with 8 and 4 respectively, suggesting more transcriptional activity was involved in structural components of the plant cells in the CONV treatment (Supplementary Figure 7).

With the overall transcript activity enrichment partitioned between structural components in the CONV treatment and the photosynthetic and organic acid cycling terms in the ORG treatment, the contigs with differential expression between tissue specific transcripts under the two treatment regimens were extracted. This was done to sort and identify the transcripts with the highest expression differences with a generalized linear model (GLM) and select the associated upper quartile slice for the GO terms (Supplementary Figure 8).

Differential expression enrichment based on GO terms in the CONV fruit tissue indicated higher representation of transcripts involved in RNA processing and oxidoreductive processes. The same four GO terms (Supplementary Figure 8) were enriched in both the GLM, and the upper quartile model. In the leaf tissue, both the GLM and upper quartile model comprised of overrepresentation of enriched GO terms for Leaf ORG and underrepresentation of non-differentially expressed contigs in the leaf CONV dataset (Supplementary Figure 9). Nine GO terms, with five of the GO terms overrepresented in the ORG leaf tissue, were enriched in the GLM model (Supplementary Figure 9A) while 13 GO terms, with six of the terms over-represented in the ORG leaf tissue, were included in the upper quartile model (Supplementary Figure 9B).

The GO terms overrepresented in the differentially expressed categories in the leaf tissues are involved in oxidoreductive and oxalate metabolic processes while the differentially expressed under-represented GO terms are involved with binding of the cyclic compounds of monosaccharides and nucleosides. Categorically, the enriched gene ontology and differential gene expression of the leaf tissue indicates more transcriptional resources are being used for photosynthetic and energetic transcript activity in the leaf CONV tissue while transcriptional resource allocation in the leaf ORG tissue was targeted towards redox maintenance activity. These observations based on GO enrichment imply that in the absence of the plant needing to divert its resources towards photorespiratory processes in CONV tissue, the metabolism is shifted towards plant growth and development, which supports part of the first hypothesis.

### Phosphate Utilization

Inorganic phosphate (P*i*) is a critical element required for plant nutrition especially for photosynthetic and respiratory metabolism. Several genes implicated in P*i* uptake are under transcriptional regulation under P*i* starvation [48]. Three of these genes; SIZ1, a SUMO E3 ligase, PHO1, a phosphate transporter and SEC12, an endoplasmic reticulum located trafficking protein, have been characterized in *Arabidopsis* as becoming upregulated during P*i* deficiency [49–54]. Expression values of *SIZ1*, *PHO1*, *SEC12* and their homologs between the ORG and CONV treatments indicate there were similar phosphate uptake profiles and any phenotypic as well as metabolite differences would not be due to phosphate deficiencies (Supplementary Figure 10).

### Impact of ORG inputs on Primary metabolic pathways

The regulation of nitrogen and carbon metabolism in plants is tightly linked. Any perturbations in nitrogen metabolism have a direct impact on the rates of photosynthesis, photorespiration, and respiration [55]. The primary source of nitrogen uptake is through the roots but the terminal sinks for nitrogen, primarily the leaf and fruit of a plant, require nitrogen influx for various metabolic processes. Since the organic fertilizer differs significantly in the type of available nitrogen, different nitrogen uptake pathways were queried for differential expression. For the analysis of this pathway, qRT-PCR based quantitative expression analysis was also included for some of the genes to complement RNAseq data and gain an understanding of how the transporters and enzymes involved in nitrogen metabolism were impacted.

#### Nitrogen Uptake – Changes in expression of nitrogen transporters

##### Nitrate transporters

Two major classes of nitrate transporters have been described in the literature. High affinity transporters operate in the 10 to 250µM nitrate concentration range and low affinity transporters operate above the 250µM concentration limit (reviewed in [56]).

Four of the nitrate transporters (**Figure 5B & C**), belonging to the NRT1-PTR family, identified in the ORG vs CONV dataset have been characterized previously as low affinity nitrate transporters [57]. All of the low nitrate affinity transporters were expressed at higher values in the leaf tissue than in the fruit tissue with three of the four exhibiting increased expression in CONV leaf compared to the ORG leaf.

**Figure 5.**
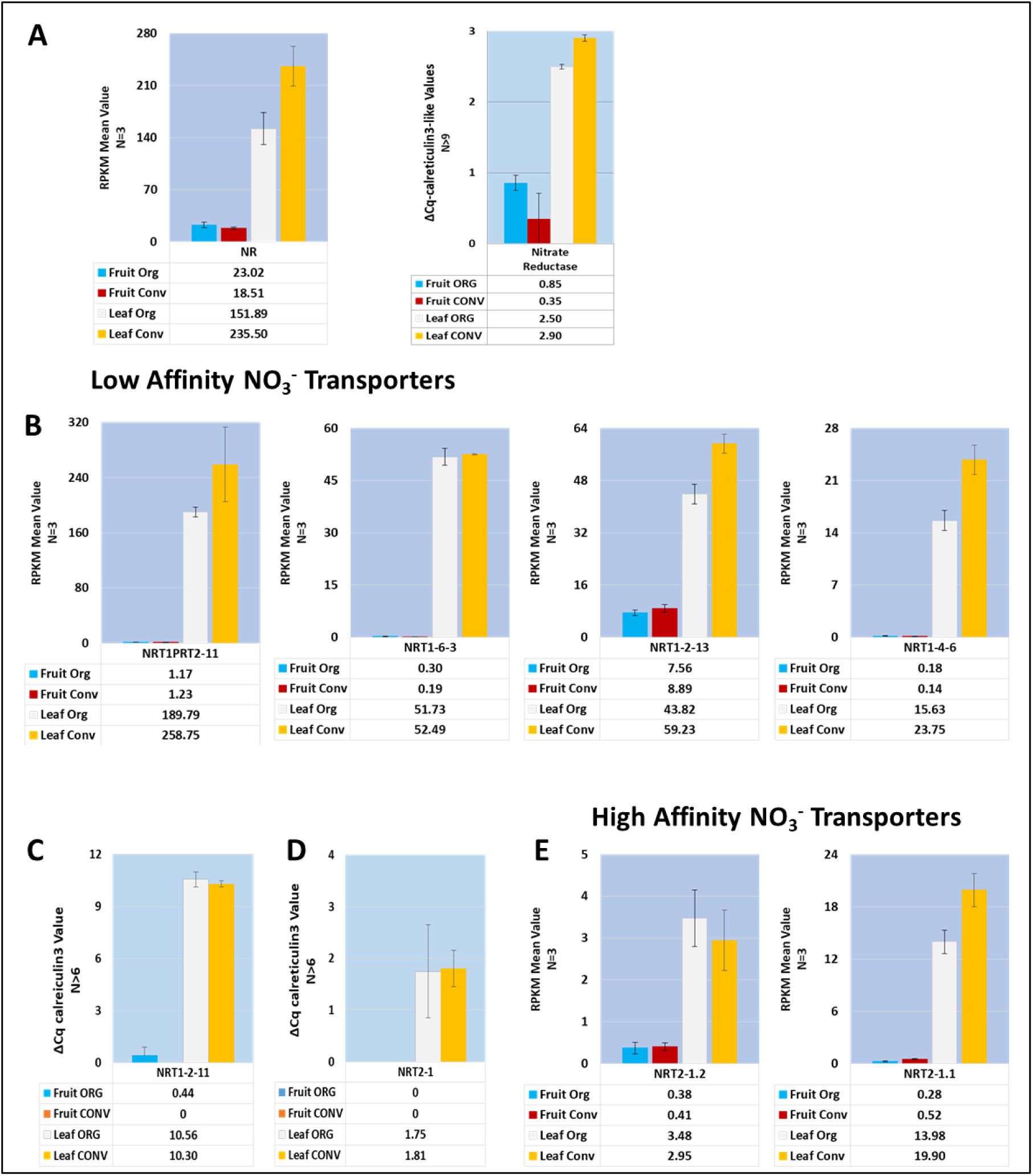
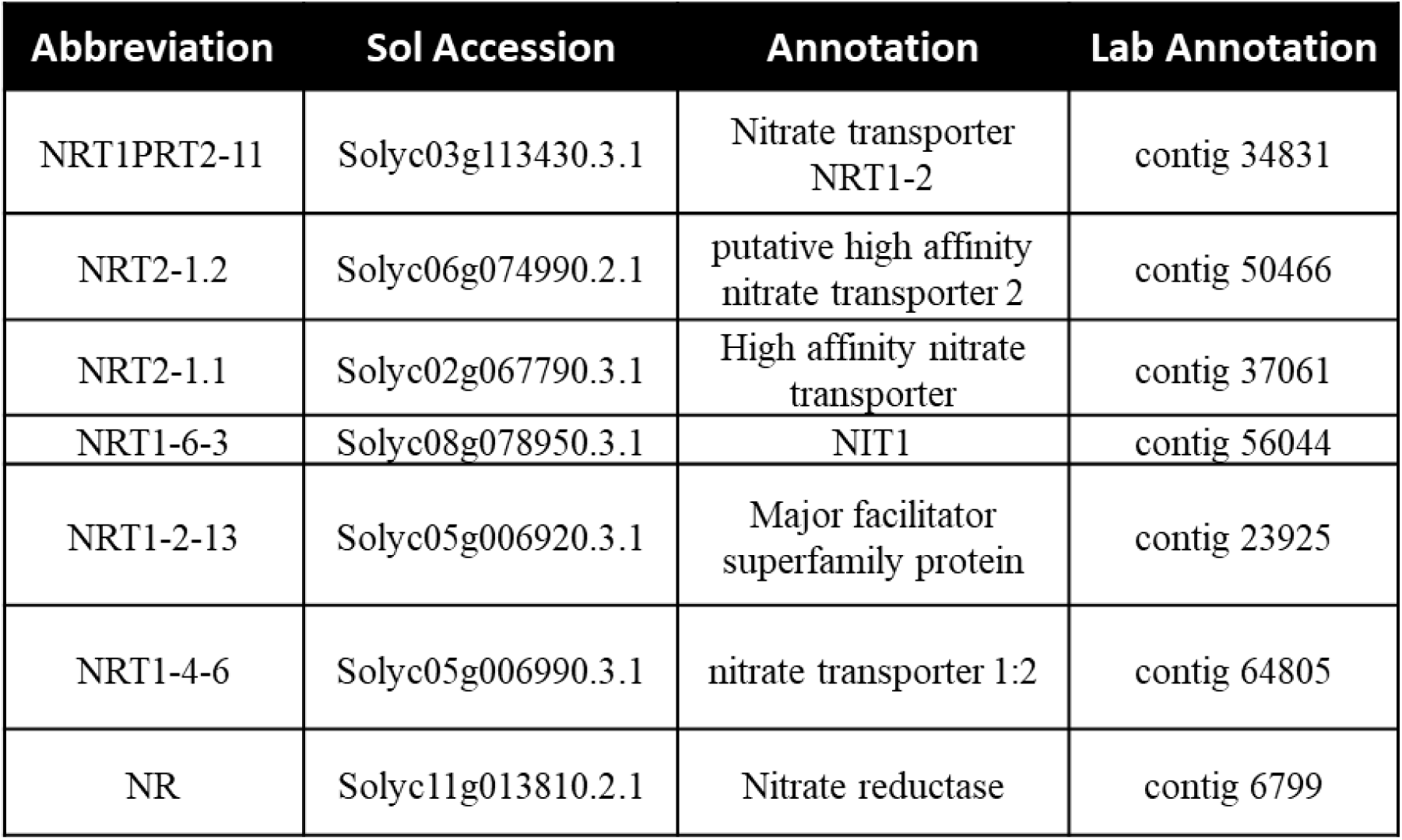
Expression profile of mean RPKM values of Nitrate Assimilation components in Fruit and Leaf tissue. **A.** RPKM and qRT-PCR values for Nitrate Reductase. **B. L**ow affinity nitrate transporters NRT1-2-11, NRT1-6-3, NRT1-2-13 and NRT1-4-6 **C & D.** qRT-PCR expression values for the highest expressed low and high affinity nitrate transporters. **E**. High affinity nitrate transporters NRT2-1.2, NRT2-1.1. qRT-PCR results (A, C and E) are averages of the number of replicates (N) and bars indicate the standard deviation between replicates. Table of annotations and accession numbers from Sol Genomics ITAG3.2. qRTPCR relative values for tissue type and treatment, trend with the corresponding RPKM values generated from the HTS analysis. Error bars denote the standard deviation of qRT-PCR expression values between 3 biological replicates and at least 3 technical replicates. Error bars denote the standard error of sequenced expression values between 3 biological replicates.

Two high affinity transporters, *NRT2-1.1* and *NRT2-1.2*, were differentially expressed between the tissue types (**Figure 5D & E**). *NRT2-1.1* was expressed in the leaf tissue with no discernable differences across the two treatments. On the other hand, *NRT2-1.2* was expressed in the Fruit ORG treatment at a level that was 30% higher than the Fruit CONV treatment. The increase of expression in the ORG treatment versus CONV treatment, points to a potential transcriptional regulation activity due to the concentration and/or bioavailability of nitrate. Nitrate deficiency in Arabidopsis has been shown to result in the up regulation of high affinity nitrate transporters [58] and correlates well with nitrate depletion in the ORG fertilizer treatment results in the fruit tissue. The low affinity transporters, belonging to the NRT1 family and involved in translocation of high molarity nitrate concentrations, while not significantly upregulated, had higher overall RPKM values, specifically with higher expression in the CONV leaf tissues.

Nitrate studies, as well as other forms of nitrogen assimilation studies, have shown nitrate, nitrite and ammonium elicits transcriptome-specific and regulatory responses in diverse and interconnected ways [59, 60]. Leaf transcript profiles from this study of the nitrate uptake and assimilation components were found to be in general agreement with the induction of transcripts from the introduction of nitrate in CONV fertilizer.

##### Nitrite transporters

Characterization of nitrite transporters remains elusive in plants as indicated by the lack of previously published literature. Given this constraint, the nitrite transporter sequences from *Arabidopsis thaliana, Cucumis sativus* and *Vitis vinifera* were used to query the *S. lycopersicum* NCBI sequence database. The XM_004240292.2 PREDICTED: *Solanum lycopersicum* protein NRT1/ PTR FAMILY 3.1-like orthologous sequence had the highest homology to the *V. vinifera* KF649633 and *C. sativus* NM_001280623 NITR1 nitrite transporters. The expression of *S. lycopersicum NITR2* orthologues were barely detectable in this study in the leaf and fruit tissues regardless of fertilizer treatment, and the expression of *S. lycopersicum* NRT1/ PTR FAMILY 3.1-like XM_004240292.2 sequence was not detected at all in the fruit tissue (data not shown). The Solyc03g113280.3.1 Integral membrane HPP family protein sequence aligned with highest homology to both the Arabidopsis NITR2.1 BT030052 and NITR2.2 BT006407 nitrite transporters and the RPKM values were found to be very low (**Figure 6C**). Thus, no conclusion could be drawn regarding the role of nitrite transporters in this experimental set up.

**Figure 6.**
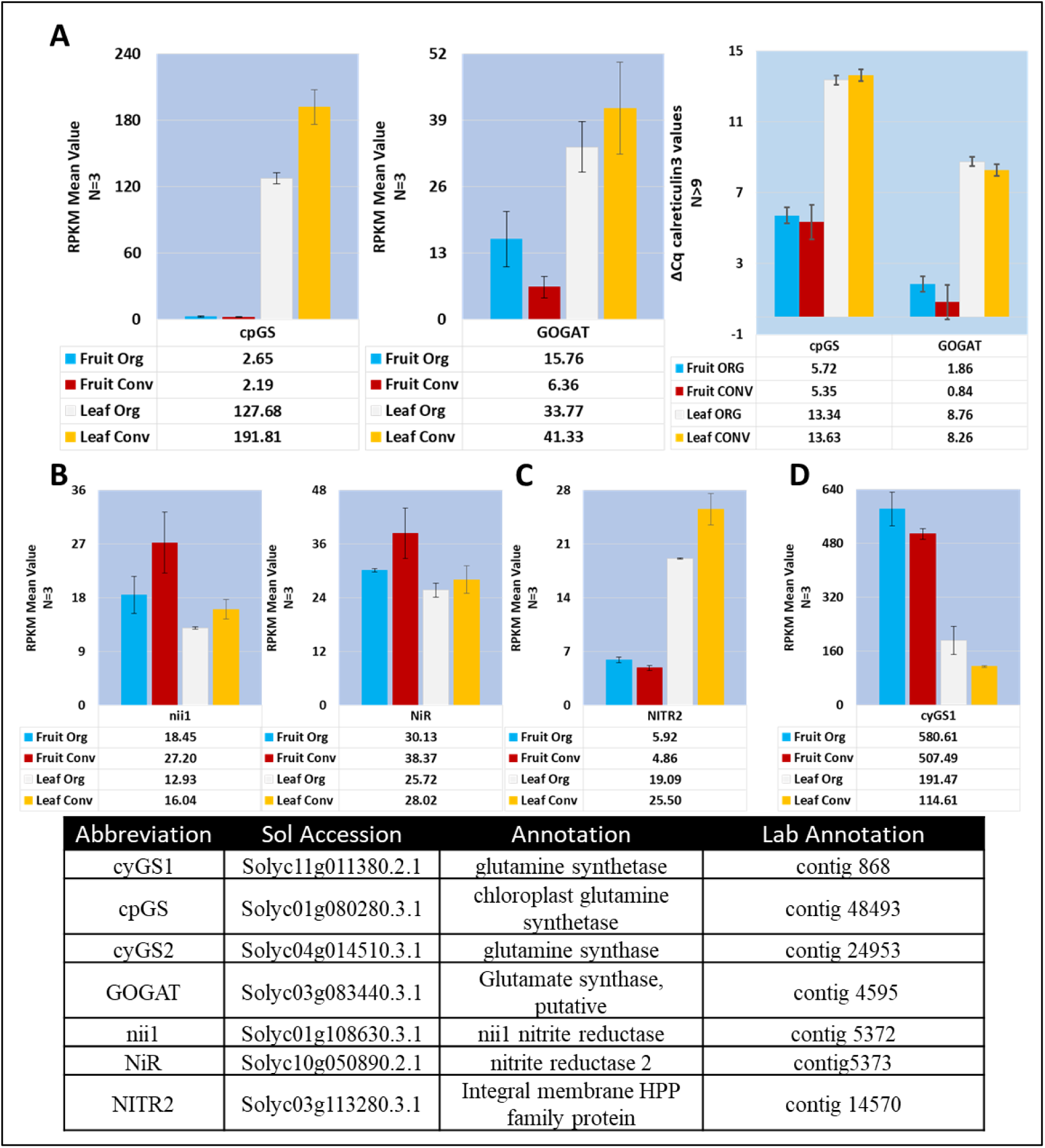
Expression profile of mean RPKM values of Nitrite Assimilation components in Fruit and Leaf tissue. **A.** Chloroplast isoform of glutamine synthetase (cpGS) and glutamate synthase (GOGAT) RPKM and ΔΔCq values. **B.** RPKM values for two isoforms of Nitrate Reductase (nii1 and NiR) **C**. Identified homolog of plastidial nitrite transporter (NITR2). **D.** RPKM values for cytosolic isoform 1 of glutamine synthetase. Table of annotations and accession numbers from Sol Genomics ITAG3.2. Error bars denote the standard error of sequenced expression values, or in the case of cpGS and GOGAT qRT-PCR ΔΔCq values, between 3 biological replicates and at least 3 technical replicates.

##### Ammonium transporters

Higher plants exhibit varied responses to ammonium (NH4^+^) and *S. lycopersicum* has been shown to be sensitive to NH4^+^ induced toxicity (reviewed in [31]. Fruit and leaf tissue produce ammonium though photorespiratory processes as well as incorporation of nitrogen from nitrate. Transcription as well as posttranslational regulation of the ammonium transporters is known to be upregulated under nitrogen deficiency [61]. Transporters implicated in ammonium translocation are categorized by the translocation fate of their primary substrate and can be found in almost every organism. The ammonium transporter/methylamine permease/rhesus (AMT1) family genes are involved in NH4^+^ uptake and NH4^+^ output and are thought to have evolved in the eukaryotes. Ammonium transporter/methylamine permease/rhesus (AMT2) genes of prokaryotic origin are involved in NH4^+^ uptake with NH3 plus a proton as the output. Three members of the AMT1 family and AMT2.1 were detected in this study. *AMT1.1*, Solyc09g090730.2.1, had the highest expression values overall in the leaf ORG tissue and leaf CONV tissue. The only ammonium transporter to exhibit higher expression values in the fruit tissue compared to leaf tissue was *AMT1.2*, Solyc04g050440.3.1, although the overall expression values were low (Supplementary Figure 11).

Conversion of toxic ammonium, whether generated through photorespiration or endogenous uptake, relies on glutamate synthase (GS) and decarboxylating glycine dehydrogenase (GDH). The *NADP-GDH isoform 2* (Solyc01g068210.3.1) was the only isoform detected at appreciable levels other than the *GDH-B* isoform (Supplementary Figure 12). While the *GDH-B* levels were slightly higher in the ORG leaf treatment and barely detected in the fruit tissue, the *NADP-GDH isoform 2* levels were higher in the fruit tissue and slightly higher in the CONV treatment when compared to the ORG tissue. It has been shown in oilseed rape that a concurrent increase in NH4^+^ flux increased GS activity [62]. The cytosolic isoform of *GS* activity was increased in the ORG treatments corresponding to the results expressed in the Schjoerring et al. 2002 (62) study (**Figure 6D**). Surprisingly, the plastidial *GS* isoform in the leaf treatment indicated the opposite trend and was barely detected in the fruit tissue (**Figure 6A**).

##### Amino Acid Transporters

Nitrogen assimilation by plants includes uptake of amino acids and plants possess transporters that are specific to different groups of amino acids [63]. In this study, homologs of transporters specific to four major groups, the proline, amino acid permease, vacuolar amino acids, and the aromatic and neutral amino acids were detected. Two proline transporter transcripts, *ProT1* (Solyc03g096390.3.1) and *ProT3* (Solyc05g052820.3.1), were identified with expression of the former being much higher than the latter, which was expressed at barely detectable levels. Neither of the proline transporters were differentially expressed between the ORG or CONV treatments and *ProT1* (Solyc03g096390.3.1) was expressed higher in the leaf tissue in comparison to the fruit tissue (Supplementary Figure 13A).

Three transcripts in the dataset were annotated as members of the amino acid permease group. *AAP3* (Solyc11g005070.2.1) exhibited highest expression in the leaf tissue with a low ORG to higher CONV trend and very low detectability in the fruit tissue. *AAPput* (Solyc07g066010.3.1), a putative homolog in the family, was expressed higher in the fruit tissues with no discernable differences between the ORG and CONV treatments and the ORG leaf tissue trended higher than the CONV tissue. *AAP2* (Solyc04g077050.3.1) expression was barely detectable in the four samples with no discernable differences between tissues or treatments (Supplementary Figure 13A).

Four members of the vacuolar transporter group were detected in this study namely, two isoforms of *AAT1*, labelled as *vacAAT1a* (Solyc10g084830.2.1) and *vacAAT1b* (Solyc11g008440.2.1), *vacCAT2* (Solyc10g081460.2.1) and *vacCAT4* (Solyc02g037510.3.1). Two additional cationic transporters, *CAT5* (Solyc02g081850.3.1) and *CAT7* (Solyc11g006710.2.1), were present but at barely detectable amounts. *vacCAT2* was the most highly expressed transporter, and was expressed higher in the fruit tissue than the leaf tissue with no discernable differences between the two treatments. *vacAAT1b* exhibited the highest differential expression between the two tissues. The fruit tissue expression values were higher than the leaf tissue with both tissues having a lower in ORG to higher in CONV trend. *vacCAT4* expression values were higher in the fruit tissue, following the trend of higher expression of this group of transporters in fruit tissue, but with no discernable differences between the ORG and CONV treatments. *vacAAT1a* expression values ran contrary to the other members of this group with barely detectable values in the fruit and very low values in the leaf tissue (Supplementary Figure 13A).

*ANT1lq* (Solyc03g117350.1.1) and the mitochondrial targeted *ANT1er* (Solyc06g069410.3.1) transcripts identified in this study belonged to the aromatic and neutral amino acid transporter group. The expression values of *ANT1er* were higher with no discernable differences between the ORG and CONV treatments but leaf tissue expression values were much higher than in the fruit tissue. *ANT1lq* expression values were very low and did not vary appreciably between tissue or treatment types (Supplementary Figure 13B). Two ureide transporters, *UPS1a* (Solyc01g010290.3.1) and *UPS1b* (Solyc04g005410.2.1), were detected with *UPS1a* expressed with the highest levels in the leaf tissue and at relatively even amounts between the treatment types in both tissues (Supplementary Figure 13B).

#### Nitrogen metabolism – Changes in expression of the enzymes

All four major enzymes involved in nitrogen metabolism, namely nitrite and nitrate reductase and, the plastidial and cytosolic isoforms of glutamine synthetase were detected in the transcriptome data. Expression of both forms of glutamine synthetase as well as nitrate reductase was assessed using quantitative reverse real time PCR (qRT-PCR).

##### Nitrate Reductase

Irrespective of the type of transporter used, nitrate is reduced to nitrite by cytosolic nitrate reductase (NR). The expression values of *NR* (Solyc11g013810.2.1) based on RPKM values were observed to be higher in the CONV leaf treatment by 29.2% in comparison to the ORG leaf treatment. Overall fruit *NR* levels were a magnitude of order lower in comparison to the leaf tissues, with the *NR* levels in ORG fruit treatment being slightly higher than the CONV fruit tissue. The expression levels of *NR*, as assayed with qRT-PCR, mirrored the expression trends observed with the RNAseq data with the CONV leaf treatment expressed at higher levels than the ORG leaf treatment and the ORG fruit treatment expression values higher than the CONV fruit treatment (**Figure 5A**).

##### Nitrite Reductase

Nitrite is transported into the plastids via the NITR family of transporters where it is converted to glutamate by ferredoxin dependent nitrite reductase. Two isoforms, *Nii1* and *NiR*, Solyc01g108630.3.1 and Solyc10g050890.2.1 respectively, of the plastid-targeted nitrite reductases were identified in the transcriptomic data. Both isoforms exhibited the same expression patterns across the different treatments and tissues. Both isoforms expressed at a higher level in the fruit tissue compared to the leaf tissue and in the CONV treatments over ORG treatments (**Figure 6B**).

#### Glutamate/Glutamine – Glutamine synthetase

Glutamate produced from nitrate reductase in the plastids is transported across the plasma membrane by the AMT1 family of transporters and converted to glutamine, in both cellular compartments, by glutamine synthetase (GS). Both compartmental forms of GS, cytosolic Solyc11g011380.2.1 (*cyGS1*) and Solyc04g014510.3.1 (*cyGS2*) and the plastidial Solyc01g080280.3.1 (*cpGS*), were identified in the transcriptome data, and two isoforms of the cytosolic GS - *cyGS1* and *cyGS2* enzyme were detected. The cytosolic form, *cyGS1*, exhibited the highest expression level in the ORG fruit tissue compared to the CONV fruit tissue as well as in the leaf tissue (**Figure 6D**). The plastidial form of *cpGS* expressed at a higher level in the CONV leaf tissue in comparison to the ORG leaf treatment and was barely detectable in the fruit tissues (**Figure 6A**). Both RPKM values and the qRT-PCR values for *cpGS* were congruent with each other with the RPKM values appearing to be more discrete (**Figure 6A**).

Increased transporter expression for nitrogen uptake is indicative of higher nitrate consumption in the leaf tissue relative to the fruit tissue as the fruit tissue transcriptional activity was scarcely detected with qRT-PCR for what appears to be a leaf specific isoform (Solyc02g067790.3.1) compared to a fruit specific isoform (Solyc06g074990.2.1) (**Figure 5C and E**). Leaf CONV tissue demonstrated a 25% increase in the expression of low affinity transporter compared to the Leaf ORG treatment across the four identified transporters (**Figure 5B**). Lower expression of the NRT1 family is indicative of lower nitrate requirements generated from downstream regulatory mechanisms. The NRT1 family of genes is known to be constitutively expressed in plants [56, 61]. Additionally, it can be inferred that the fruit tissue nitrate requirements were low from the lack of transcriptional activity exhibited by the NRT1 family of transporters. As nitrate is not utilized by the plant directly, but incorporated as ammonium, transcriptional activity, associated with the enzymes and transporters from nitrate to nitrite to ammonium, was higher in the leaf CONV treatments with the exception of the nitrite reductase isoforms expressed at marginally higher levels in the fruit CONV tissue and treatment and a highly expressed cytosolic glutamine synthetase isoform in the fruit ORG tissue (Figures 5 & 6). Leaf ammonium and glutamine levels have been shown to have a negative correlation to nitrate reductase transcription activity [59]. Transcript levels of nitrate reductase in the leaf ORG and both fruit tissue treatments implicate higher levels of ammonium, glutamate and glutamine in these tissues. The transcript activity differential between glutamine synthetase and glutamate dehydrogenase suggest the metabolic flux runs in favor of the production of glutamate. Additionally, the increase of glutamate would add to the increase of umami flavor, as well as the significant increase in soluble solids that contribute to consumer taste preference of tomatoes [64].

#### Photosynthetic/Respiratory Processes

The Gene Ontology enrichment of the transcriptome data revealed that the photosynthetic components and processes, which include photorespiratory and respiratory components and processes, were enriched in the ORG fruit tissues while the translational machinery and nonspecific organelle components were enriched in the CONV fruit tissue. This may be somewhat surprising since mature fruit tissues are not expected to perform any photosynthesis. However, GO terms implicated in photosynthesis are included in the photorespiratory and respiration processes. Analysis of the annotations of the enriched GO terms revealed that enzymes involved in the TCA cycle, as well as the C4 photosynthetic cycle, were enriched while the expression of transcripts coding for chlorophyll binding protein was reduced in the ORG treatment.

Genes related to the plant’s photosynthetic processes; the ATP-dependent zinc metalloprotease FtsH in chloroplast development [65], photosystem I subunit B (psaB) and photosystem II subunit A (psbA) related to the electron transport chain [66], ribulose-1,5-bisphosphate carboxylase/oxygenase (rubisco) large subunit for carbon/oxygen fixation [67], peroxisomal (S)-2-hydroxy-acid oxidase (glycolate oxidase, GLO) involved in remediation of the fixed oxygen via photorespiration [68], and the superoxide dismutase, L-ascorbate peroxidase (APX) and catalase genes [69] involved in the scavenging of reactive oxygen species are useful in understanding the photosynthetic status of the plant. RPKM expression values related to genes involved in the capture and conversion of energy (Supplementary Figure 14) were indicative of more transcriptional activity in the CONV leaf treatment. Conversely, *APX.1* and *SOD Cu-Zn.1*, the genes related to photosynthetic stress remediation of ROS scavenging (**Figure 3**) demonstrated an increased trend in ORG leaf and ORG fruit tissue, which imply the ORG treated plants were under photosynthetic stress. The antioxidative metabolites produced during photosynthetic stress are known to be photosynthetic feedback regulators and known to inhibit the expression of photosynthetic genes.

The TCA cycle is known to either operate in a closed mode where the cycle functions normally with all the enzymes operating to produce chemical energy and amino acid precursors, or in an open mode, leading to the accumulation of different organic acids dependent on where the flux break occurs. These breaks can occur due to changes in redox status or under photorespiratory conditions (Igamberdiev and Eprintsev, 2016). The alterations in gene expression networks may be responsible for allocation of photosynthetically derived resources to the enhanced synthesis of phytonutrients observed in ORG fruit treatment.

The enzymes involved in the photosynthetic C4-cycle were represented by contigs annotated as Phosphoenolpyruvate carboxykinase (PEPCK), phosphoenolpyruvate carboxylase (PPC), malate dehydrogenase (MDH), Pyruvate Kinase (PyK) and NADP-ME. These enzymes are important for the conversion between the organic acids for the precursors of amino acids, production of lipids, generation of Krebs cycle precursors, production of sugars and NADPH during respiration as the fruit ripens.

Phosphoenolpyruvate carboxykinase (PEPCK) is, *in vitro*, a readily reversible enzyme, responsible for the conversion of oxaloacetic acid (OAA) to phosphoenolpyruvate (PEP) or, in the reverse direction, PEP to OAA depending on the flux pressure of the intermediates (Walker et al., 2016). Five isoforms of *PEPCK* were identified in the transcriptome data with *PEPCK ATP dependent* (Solyc04g076880.3.1) and *PEPCK 2.1* (Solyc06g053620.3.1) isoforms exhibiting 3-fold and 10-fold expression in the fruit tissue respectively (Supplementary Figure 15A). The other isoforms, *PEPCK1.1* (Solyc04g009900.3.1) and *PEPCK1.2* (Solyc04g009910.3.1) exhibited higher expression in the leaf tissues where *PEPCK2.2* (Solyc09g090090.2.1) was barely detected. The overall expression was not different between the treatments.

In the context of ripening and respiration, the malic enzymes (ME) and family of malate dehydrogenases (MDH) are responsible for the decarboxylation and/or oxidation of accumulated malate pools to pyruvate and OAA respectively (Supplementary Figure 16B & C). In this transcriptome data, seven *MDH* isoforms, targeted to different cellular locations; cytosol – *MDH1* (Solyc09g090140.3.1) and *MDH-like 2.1* (Solyc09g091070.2.1) and *MDH-like 2.2* (Solyc03g115990.2.1), glyoxysome – *glyMDH3.1* (Solyc02g063490.3.1) and *glyMDH3.2* (Solyc01g106470.3.1), and chloroplast – *cpNADPMDH1* (Solyc11g007990.2.1) and *cpNADPMDH 2* (Solyc03g071590.3.1), were identified. The *MDH1* transcripts were present at the highest levels in the leaf tissue and the *glyMDH3.1* isoform in the fruit tissue compared to the other *MDH* isoforms. The *MDH1* and *glyMDH3.1* transcripts were highly abundant in fruit tissues although there were only slight differences between the CONV and ORG treatments (**Figure 7** and Supplementary Figure 16C).

**Figure 7.**
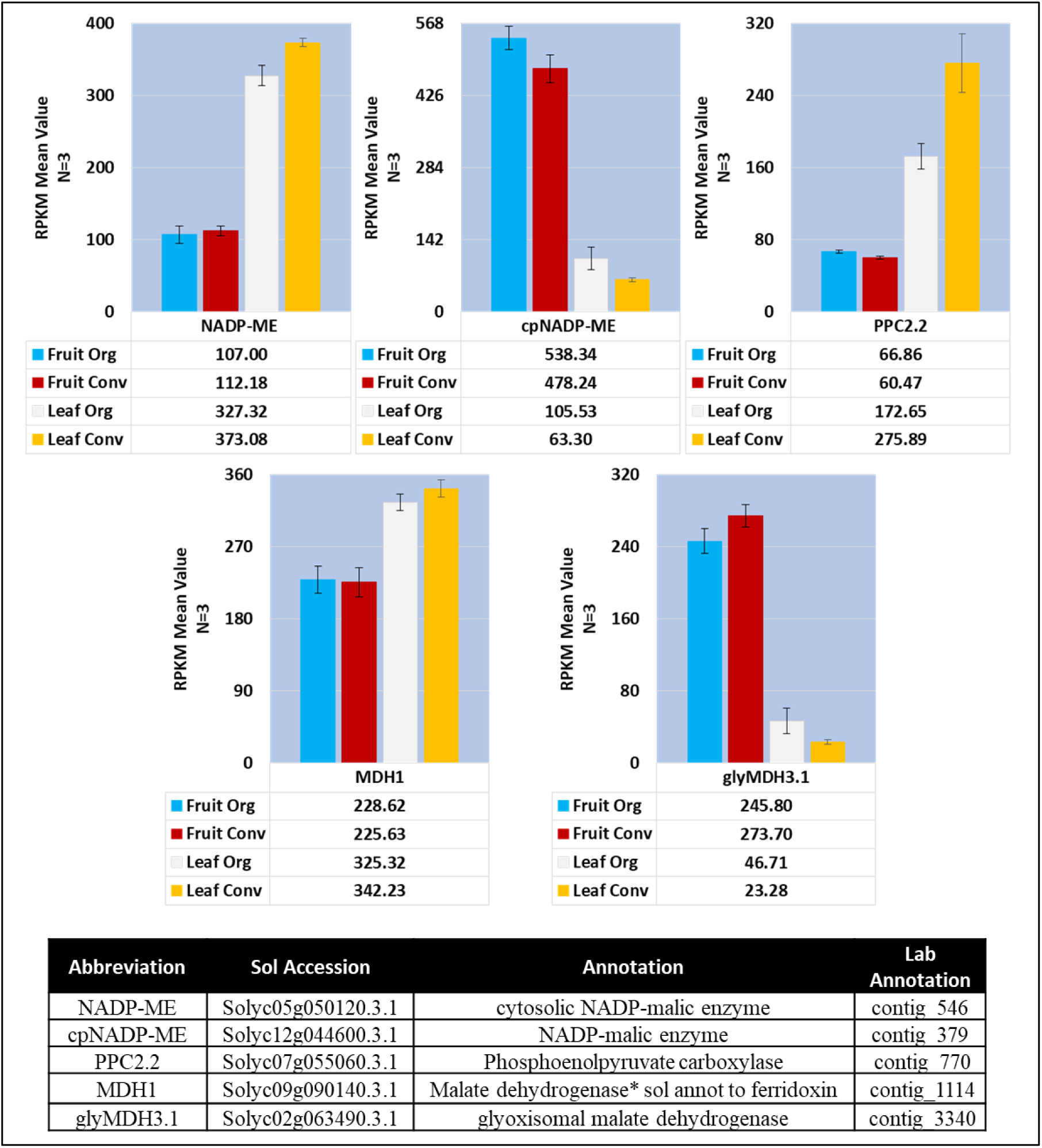
Mean RPKM expression values of enzymes involved in photosynthetic and/or tricarboxylic acid cycle respiratory pathway genes. Table of annotations and accession numbers from Sol Genomics ITAG3.2. Error bars denote the standard error of sequenced expression values between 3 biological replicates.

Five different isoforms of phosphoenolpyruvate carboxylase (*PPC*) were identified; two isoforms of *PPC1*, two isoforms of *PPC2* and a *PPC4* isoform.

The *PPC2* isoform (Solyc07g055060.3.1), had the highest expression values for both the fruit and leaf tissues. The leaf tissue CONV expression value was 35.5% higher than the ORG expression value (Supplementary Figure 16B). The expression values of the second *PPC2* isoform (Solyc07g062530.3.1) were highest in the fruit tissue. Leaf tissue values were markedly lower for the ORG and CONV treatments. The *PPC1.1* (Solyc12g014250.2.1) and *PPC1.2* (Solyc10g007290.3.1) contigs had comparable expression in the leaf ORG and CONV dataset. The fruit tissue expression pattern for *PPC1.2* was almost nonexistent and *PPC1.1* had values of 11.86 in the fruit ORG and 15.00 in the fruit CONV treatments. The lone *PPC4* isoform (Solyc04g006970.3.1) was expressed at a low level across both tissue types (Supplementary Figure 16B).

There were seven pyruvate kinase contigs identified in the *de novo* transcriptome. Four cytosolic and 3 chloroplast isoforms were detectable in both the fruit and leaf tissue types across the two treatments, however one cytosolic gene, *PyKfp2* (Solyc10g083720.2.1), and one chloroplast-targeted gene, *PyK1.2* (Solyc09g008840.3.1) had notable expression levels (Supplementary Figure 15B). Both genes were expressed at a higher level in the fruit tissues with the *PyKfp2* expressing at over 3.5-fold higher level compared to the *PyK1.2*. There was no appreciable difference in expression in either isoform across treatments.

NADP-Malic Enzyme (NADP-ME) was represented by four genes, with three cytosolic (NADP-ME and NADP-ME1.1 and 1.2) and one targeted to the chloroplast (cpNADP-ME)(Supplementary Figure 16A). Similar to the expression pattern of other genes, the expression of all the family members was not appreciably different across treatments. However, *NADP-ME1* was predominantly expressed in the leaf tissues where *NADP-ME1.1* and *cpNADP-ME* were predominantly expressed in the fruit tissues (**Figure 7**).

Expression of chlorophyll a/b binding protein (CAB) transcripts in this study, while not differentially expressed between the treatments, exhibited a reduced overall expression in ORG leaf (Supplementary Figure 17). In addition, a differential increase in the expression of *APX* and *cpSOD Cu-Zn* in the ORG leaf was observed (**Figure 3**). In addition to the increase in these genes, the decrease in photosystem I and photosystem II reaction center transcripts, *psaB* and *psbA* (Supplementary Figure 14) is an additional indicator that the ORG treated plants did not hold as much capacity for photosynthetic activity as the CONV treated plants did. These observations indicate that the plants under the ORG treatment were operating under photosynthetic stress conditions where a reduction in chlorophyll content and increase in the reactive oxygen species (ROS) scavenging genes could be used as mechanisms to reduce production of ROS. Under such conditions, plants attempt to decrease photosystem II activity by activating the Mehler peroxidase reaction (reviewed in [70], which results in an increase in superoxide radicals. Increased oxygen radicals stimulate SOD production [71, 72], and APX and catalase expression activity during tomato fruit ripening [73]. Increased level of APX expression was also observed in *Lactuca sativa* L. cv Romaine leaves when Mehler reaction was initiated by potassium cyanide (KCN)-mediated inhibition of the electron transport chain [74]. In this study, downregulation of ORG leaf *CAB*s, differential increase of the *APX* and *cpSOD Cu-Zn* isoforms in ORG leaf and upregulation of *Psy1* in ORG leaf discussed previously imply that ORG treatment produces a photoinhibitory environment for the plant.

## CONCLUSIONS

In this study two hypotheses were evaluated. The first hypothesis that organic fertilizer, which results in a slower rate of biological release of available nitrogen to plant roots, results in greater allocation of photosynthetically derived resources to the synthesis of secondary metabolites, such as phenolics and other antioxidants, than to plant growth, was supported by the higher accumulation of lycopene, ascorbate and TEAC.

The second hypothesis stated that the genes involved in changes in the accumulation of phytonutrients under organic fertilizer regime will exhibit differential expression, and that the growth under different fertilizer treatments will elicit a differential response from the tomato genome, was partially supported. Plant growth under different fertilizer treatments elicited a differential response from the tomato genome (Figure 1, Supplementary Figure 3A and B). While the observed expression pattern of genes involved in metabolic pathways of interest did not always follow expected expression trends, pathway analysis provided critical information regarding the metabolic state of the plants grown under ORG fertilizer treatment. The CONV enriched GO categories in the fruit of the cellular macromolecule metabolic process, which include the RNA metabolic process and the cell wall macromolecule process correlate well with the findings of [75].

Based on the metabolic pathway modeling presented in this study, we propose that the organic fertilizer treatment results in activation of photoinhibitory processes through differential activation of nitrogen transport and assimilation genes resulting in higher accumulation of the phytonutrients quantified in the fruit tissues. These data also provide information regarding which genes are impacted the most due to the differences in source of fertilizer. This knowledge can be used to identify alleles that allow for efficient utilization of organic inputs for breeding crops that are acclimatized to organic inputs. From this analysis it is also clear that in order to gain a deeper understanding of the metabolic flux under the ORG treatment, it is essential to perform a time course developmental transcriptome and metabolome analysis of the fruit, leaf and root tissues.

## MATERIALS AND METHODS

### Plant material

Tomato seeds (*Solanum lycopersicum* L.) ‘Oregon Spring’ sourced from Johnny’s Selected Seeds, Winslow, ME, were sown in LC1 Professional Growing Mix (Sun Grow Horticulture, Bellevue, WA). Glasshouse temperatures were maintained at 21.1/18.3°C (day/night) with a 14 h day, supplemented with high-pressure sodium lamps and 10h night photoperiod. Three-week emergent seedlings were fertilized with a BioLink All Purpose Fertilizer 5-5-5 (Westbridge Agricultural Products, Vista, CA) solution at a concentration of 4 mL/L tap water representing the organic nutrient treatment (ORG). The seedlings grown under conventional fertilizer (CONV) received a Peters 20-10-20 solution (1.02 g/L tap water). Both treatments received one liter per week of their respective nutrient treatment.

Six-weeks-old plants of similar size and vigor from each treatment were transferred to individual 24-liter pots. Potting media for the ORG nutrient treatment consisted of a mix of LC1 Professional Growing Mix (Sun Grow Horticulture, Bellevue, WA), Whitney Farms Compost (Scotts, Marysville, OH), and sifted soil in a ratio by volume of 15:5:1. The CONV plants were potted in 100 percent LC1 Professional Growing Mix. Upon transplantation, individual plants were fertilized with 150 mL of applicable nutrient solution once per week. Twelve plants in each nutrient treatment were selected at week seven and arranged in a randomized complete block design consisting of six blocks. Plants were provided 1 liter of water every other day and the dosage of weekly nutrient treatments was increased to 500 mL beginning at week eight. Lateral shoots below the first flower cluster were removed as they appeared. Starting in week 11, the CONV nutrient solution concentration was increased from 1.02 g/L of Peters 20-10-20 to 1.25 g/L. Beginning in week 12, nutrient treatments were augmented with micronutrients. The ORG treatment was amended with BioLink Micronutrient Fertilizer (Westbridge Agricultural Products, Vista, CA) at 3.9 mL/L tap water and the CONV treatment was augmented with calcium phosphate monobasic monohydrate at 166 mg/L tap water. Concentrations of macronutrients were equivalent in both treatments, with total nitrogen, total phosphorus and total potassium concentrations of 260 ppm, 99 ppm and 235/220 ppm (CONV/ORG respectively) based on laboratory analyses (Analytical Science Laboratory, University of Idaho, Moscow, ID). Nitrogen forms varied greatly between CONV and ORG treatments with nitrate constituting the dominant form in CONV, and organic forms (i.e. amino acids and proteins) predominating the ORG nutrient solution. In week 13, the air temperature in the glasshouse was increased to 23.3/20°C (day/night) and the nutrient dosage was increased to 750 mL per plant every other day. In week 14 the nutrient dosage was increased to 1 liter every other day.

Fruit were harvested at ripe red stage beginning in week 18 and continuing until a minimum of eight ripe fruit were harvested from each plant during week 23. Immediately following harvest, all tomato fruit were weighed and their stages of maturity noted based on previously established standards [76]. A 1 cm thick slice was cut from the equatorial region of each fruit and the pericarp tissue external to each locule was sampled. The tissue was minced with a knife, one portion of which was used to measure percent soluble solids (°Brix) using a digital refractometer (Atago PR-101, Bellevue, WA), while the remainder was flash-frozen with liquid nitrogen, placed in 50-mL plastic tubes, and stored in a −80°C freezer for biochemical and transcriptome analysis. The samples were ground by mortar and pestle in liquid nitrogen until finely powdered and used for phytochemical analyses. Analyses of total carbon and total nitrogen (via combustion) and calcium, potassium, magnesium, sodium, phosphorus and sulfur (ICP with nitric digestion) were conducted on dried leaf tissues (Analytical Science Laboratory, University of Idaho, Moscow, ID).

Total phenolic compounds were measured spectrophotometrically using the method outlined by Singleton et al. (1999) [47] with modifications. One milliliter of an 80% methanol solution was added to frozen, 350 mg samples of powdered fruit tissue or 150 mg of powdered leaf tissue in 2 mL microcentrifuge tubes, with all samples run in triplicate. Tubes were vortexed for 30 sec until thoroughly dispersed and stored at −20°C for 24 hr. Samples were centrifuged at 8,000 g and 4°C for 20 min. The supernatant from each tube was then poured into individual 15 mL polyethylene sample tubes, capped and stored at −20°C. The above extraction was conducted two more times on the same tissue. The supernatant was adjusted to 3 mL with 80% methanol. Then, 200 μL of extract, 1000 μL of 10% Folin-Ciocalteu (F-C) phenol reagent, 2N and 800 μL of 75 g L-1 Na2CO3 were pipetted into 2 mL microcentrifuge tubes, vortexed and stored at 20°C for 2 hr. A second solution with each extract was made substituting 800 μL of Nanopure water for the Na2CO3 and stored at 20°C for 2 hr. An ultraviolet-visible spectrophotometer (Agilent Technologies, Model HP8453, Avondale, PA) interfaced to a computer with UV-Visible ChemStation software (v. B.01.01) with tungsten bulb was blanked on 1.0 mL of 10% F-C reagent in a 1.5 mL polystyrene cuvette. One milliliter of the sample solutions was pipetted into separate cuvettes and absorbance was measured at 760 nm. Cuvettes were run in duplicate. The average difference in absorbance was compared to a standard curved for gallic acid, and the concentrations were reported as gallic acid equivalents (GAE).

Lycopene was measured spectrophotometrically using the method outlined by Nagata and Yamashita (1992) [77] with modifications. The following work was conducted under low light. Powdered, frozen fruit samples of 100 mg were placed in 15 mL conical polyethylene sample tubes wrapped with aluminum foil with all samples run in triplicate. To each tube, 8.33 mL of a 2:3 solution of HPLC-grade acetone and HPLC-grade hexanes at 4°C was added. The solution was amended with 9.08 × 10-4 mol L-1 (200 mg L-1) butylated hydroxytoulene (BHT) to prevent pigment oxidation. Tubes were capped and vortexed for 1 min, then stored at −20°C for 24 hr. After storage the tubes were again vortexed for 1 min and returned to −20°C. This was continued for a total of 96 hr. of extraction. Tubes were vortexed a final time and let stand for 5 min for a complete phase separation to occur. The spectrophotometer was blanked and zeroed using a tungsten lamp and 1 mL 100% HPLC-grade hexanes in a 1.4 mL glass cuvette at 505 nm. Aliquots of 1 mL of the hexane phase were pipette into 1.4 mL glass cuvettes and absorbance was measured at 505 nm. Lycopene concentration was calculated according to the equation of Davies (1976): Lycopene (mM) = Absorbance505/3400 mM −1

For transcriptome analysis, samples were processed via pulverization under liquid nitrogen in the SPEX SamplePrep® FreezerMill 6870 (Metuchen, NJ USA). Three to four randomly selected representative leaves were harvested at week 23 as sub-samples from the mid to upper canopy of all plants, flash-frozen with liquid nitrogen and stored in a −80°C freezer and pulverized in the SPEX SamplePrep® FreezerMill 6870 (Metuchen, NJ USA) prior to extraction of RNA.

### RNA extraction

RNA was isolated from CONV and ORG tomato fruit and leaf tissue at the red ripe stage with the QIAGEN RNeasy Plant Mini Kit (Valencia, CA). Extractions were performed by combining three to four fully expanded leaves from the mid to upper canopy (leaf samples) or three red ripe fruit as they became ripe (fruit samples) from two plants constituting one biological replicate. RNA was extracted from each biological replicate; three CONV fruit, three ORG fruit, three CONV leaf and three ORG leaf were used as input for sequencing and qRT-PCR procedures. RNA concentration was determined for each extraction of each unique treatment-sample combination using a Nanodrop ND-8000 (ThermoFisher, MA, USA) and sample integrity was validated using denaturing agarose gel electrophoresis. The samples were treated with DNAse using the Ambion TURBO DNA-free kit (ThermoFisher, MA, USA) to remove any potential DNA contamination. RNA integrity and concentration were verified for all samples following DNAse treatment.

### Illumina Sequencing

RNA extractions were used to generate twelve individually barcoded sequencing libraries with Illumina’s TruSeq RNA Sample Preparation v2 kit (San Diego, CA, USA) with minor modifications. Modifications to the published protocol include a decrease in the mRNA fragmentation incubation time from 8 minutes to 30 seconds to create the final library proper molecule size range. Additionally, Aline Biosciences’ (Woburn, MA, USA) DNA SizeSelector-I bead-based size selection system was utilized to target final library molecules for a mean size of 450 base pairs. The quantity and quality of all the libraries were then ascertained using Life Technologies (Carlsbad, CA, USA) Qubit Fluorometer and an Agilent (Santa Clara, CA, USA) 2100 Bioanalyzer (Dr. Jeff Landgraf, Michigan State University, personal communication). The Illumina Hi Seq 2000 sequencing platform (San Diego, CA, USA) was used to sequence the cDNA libraries as 2×100 PE reads across three lanes of a flowcell at Michigan State University’s Research Technology Support Facility. Read files were submitted to the National Center for Biotechnology Information (NCBI) Short Read Archive (SRA) database under SRR4102059 (leaf conventional treatment), SRR4102061 (leaf organic treatment), SRR4102063 (fruit conventional treatment), and SRR4102065 (fruit organic treatment).

### Data Processing, Assembly, identification of differentially expressed genes and visualization of Genome-wide expression

Sequence read information from Illumina HiSeq 2000 2×100 PE fastq files were used as input for the CLC Bio Genomic Workbench (ver 6.0.1)(Aarhus, Denmark). All read datasets were processed with the CLC Create Sequencing QC Report tool to assess read quality. The CLC Trim Sequence process was used to trim the first 12 bases due to GC ratio variability and for a Phred score of 30. All read datasets were trimmed of ambiguous bases. Illumina reads were then processed through the CLC Merge Overlapping Pairs tool and all reads were *de novo* assembled to produce contiguous sequences (contigs). Mapped reads were used to update the contigs and contigs with no mapped reads were ignored. Non-trimmed reads used for assembly were mapped back to the assembled contigs. Consensus contig sequences were extracted as a multi-fasta file. The individual CONV and ORG read datasets, original non-trimmed reads, were mapped back to the assembled contigs to generate individual treatment sample reads per contig and then normalized with the Reads Per Kilobase per Million reads (RPKM) method [78].

The ORG/CONV RPKM value ratios, calculated from loci mapped on the *S. lycopersicum* genome build 2.40 chromosomal maps provided by the Sol Genomics Network [79], obtained from the RNAseq data analysis were log10 transformed and the comparison between ORG and CONV treatments for each discrete area on the chromosome was visualized via Manhattan plots (Figure 1, Supplementary Figure 3A and B). At any given position on the chromosome, red bars indicate a higher log10 ratio in the CONV treatment, and blue bars indicate higher expression in the ORG treatment. Trend lines were graphed to indicate a predominance of chromosomal activity either in the ORG treatment (positive y-intercept value) or in the CONV treatment (negative y-intercept value).

### Functional Annotation

Assembled contiguous sequences (contigs) were annotated by alignment with blastx through Blast2GO [22] (BioBam Bioinformatics S.L., Valencia, Spain) as well as local stand-alone blastx alignments against the NCBI nr database (ver. 2.2.27+) [80] and blastn alignments against the ITAG3.2_CDS.fasta file downloaded from the Sol Genomics Network FTP site (ftp://ftp.solgenomics.net/genomes/Solanum_lycopersicum/annotation/ITAG3.2_release/). Gene ontology (GO) annotation, enzyme code annotation and the EMBL-EBI InterProScan annotation of predicted protein signatures were all annotated through Blast2GO [21, 22]. The BLAST annotated RNA-Seq datasets from the conventional and organic treatments were analyzed for GO enrichment with Blast2GO [21]. Unless otherwise specified, expression analysis was restricted to the contig consensus sequence annotation and does not represent specific alleles, gene family members of highly similar sequence or subunit specificity [81].

Fisher’s exact test was performed to identify any nonrandom associations between the Gene Ontology terms associated with the annotated transcripts to identify any over- or under-represented GO terms involved in the phytonutrient pathways related to TEAC, total phenolics, and soluble solids. ORG/CONV RPKM ratios were log10 converted and separated as 2-fold greater, log10 value less than −0.3010 – higher in the CONV treatment or log10 greater than 0.3010 – higher in the ORG treatment, between the treatments and non-differential, log10 values between −0.3010 and 0.3010 (Supplementary Figure 8 and 9). Contig read counts were included in the Blast2GO analysis and queried with Fisher’s exact test for differential expression within a General Linearized Model (GLM), and the upper quartile between ORG and CONV leaf and fruit treatments (Supplementary Figure 8 and 9).

### Quantitative RT-PCR

RNA extractions used to generate the twelve individually barcoded sequencing libraries, one from each biological replicate, were utilized for first-strand cDNA synthesis for subsequent quantitative polymerase chain reactions using the Invitrogen SuperScript VILO kit in 20μl reactions. Product integrity was checked using agarose gel electrophoresis. Concentration for each cDNA preparation was evaluated using a Qubit fluorimeter (Life Technologies – Carlsbad, CA, USA). Quantitative real-time PCR (qRT-PCR) was performed in triplicate from each unique cDNA preparation derived from three independent experiments. The abundance of 19 differentially expressed genes selected from *in silico* RPKM analysis of RNAseq data was quantified. Each reaction was loaded with 25 ng of template first-strand cDNA, and tested on the Stratagene MX3005P light cycler (Agilent Technologies – Santa Clara, CA) using the iTaq Universal SYBR Green Supermix reagent (with ROX passive dye) (Bio-Rad Laboratories – Hercules, CA). Reaction conditions and thermal profile can be found in the recommended protocols provided by the manufacturer. Instrument fluorescence was used as an input for the LinRegPCR tool [82, 83] and analyzed using the platewide-mean mode for extraction of reaction Cq and efficiency values. Fold-change values of transcript abundance were calculated relative to the geometric mean of internal control of calreticulin3-like transcripts and represented using the Pfaffl-method correction [84].

## Acknowledgments

This research was funded in part by CSANR BIOAg grant to PA and AD, and by the USDA National Institute of Food and Agriculture, Hatch project WNP00011 to AD. SLH and BRK acknowledge the support of NIH/NIGMS through institutional training grant award T32-GM008336. The contents of the publication are solely the responsibility of the authors and do not necessarily represent the official views of the NIGMS or NIH.

## Legends to Supplementary Tables and Figures

Supplementary Table 1: Above and below ground vegetative biomass on fresh (FW) and dry weight (DW) bases, and percent root biomass fraction for conventional (CONV) and organic (ORG) fertilizer treatments

Supplementary Table 2: Mineral concentration of leaves on a dry weight (DW) basis for conventional (CONV) and organic (ORG)

Supplementary Table 3. Sequenced and assembled reads.

Supplementary Table 4. Sol Genomics Network ITAG ver. 3.2 CDS annotations to gene lab abbreviations from Supplementary Figure 5.

Supplementary Table 5. Sol Genomics Network ITAG ver. 3.2 CDS annotations to gene lab abbreviations from Supplementary Figure 6a and 6b.

Supplementary Figure 1: Mean red ripe fruit mass under conventional (CONV) and organic (ORG) fertilizer treatments

Supplementary Figure 2. qRT-PCR reference gene validation across tissue and treatment types.

Supplementary Figure 3A. ORG and CONV log10 RPKM ratio Manhattan plot representations of expression values across S. lycopersicum chromosomes 1 to 6.

Supplementary Figure 3B. ORG and CONV log10 RPKM ratio Manhattan plot representations of expression values across S. lycopersicum chromosomes 7 to 12.

Supplementary Figure 4. Lycopene Biosynthesis Pathway Enzymes.

Supplementary Figure 5. Mean expression values of the Smirnoff-Wheeler pathway enzymatic transcript activity.

Supplementary Figure 6a. Mean expression values of the Foyer-Halliwell-Asada Pathway enzymatic transcript activity

Supplementary Figure 6b. Mean expression values of the Foyer-Halliwell-Asada Pathway enzymatic transcript activity.

Supplementary Figure 7. Differential expression breakout of contigs assigned to the enriched GO terms.

Supplementary Figure 8. Fruit GO terms concluded to be differentially expressed by the general linear model (A) and for the upper quartile (B).

Supplementary Figure 9. Leaf GO terms concluded to be differentially expressed by the general linear model (A) and for the upper quartile (B).

Supplementary Figure 10. Mean expression values of the Phosphate uptake indicator genes.

Supplementary Figure 11. Mean expression values of Ammonium Assimilation transporters in Fruit and Leaf tissue.

Supplementary Figure 12. Mean expression values of the Glutamate Dehydrogenase genes

Supplementary Figure 13. Detected and identified amino acid transporters and mean expression values

Supplementary Figure 14. Mean expression values of Photosynthesis Indicator genes.

Supplementary Figure 15. Differential expression of Phosphoenolpyruvate carboxykinase and Pyruvate kinase genes.

Supplementary Figure 16. Photosynthetic C4 and/or Respiratory pathway genes.

Supplementary Figure 17. Chlorophyll A/B binding proteins and RPKM mean expression values.

