## Supplementary figures and images for "Concomitant phytonutrient and transcriptome analysis of mature fruit and leaf tissues of tomato (*Solanum lycopersicum* L. cv. Oregon Spring) grown using organic and conventional fertilizer"

### Supplementary Figure 1

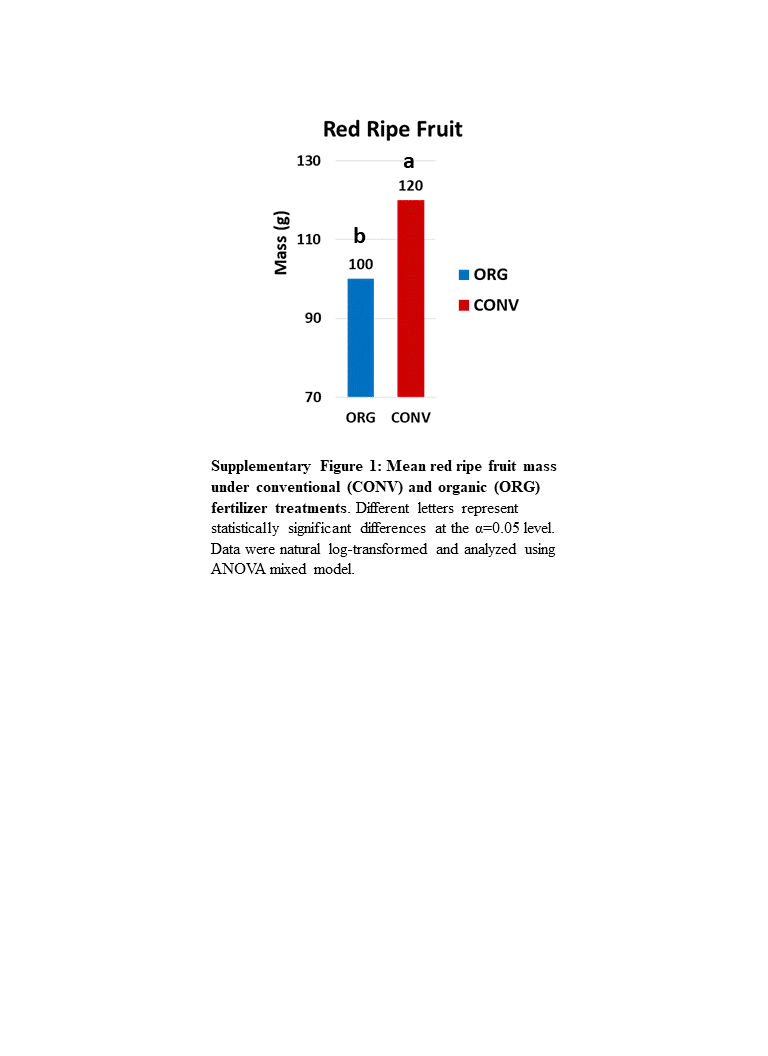

### Supplementary Figure 2

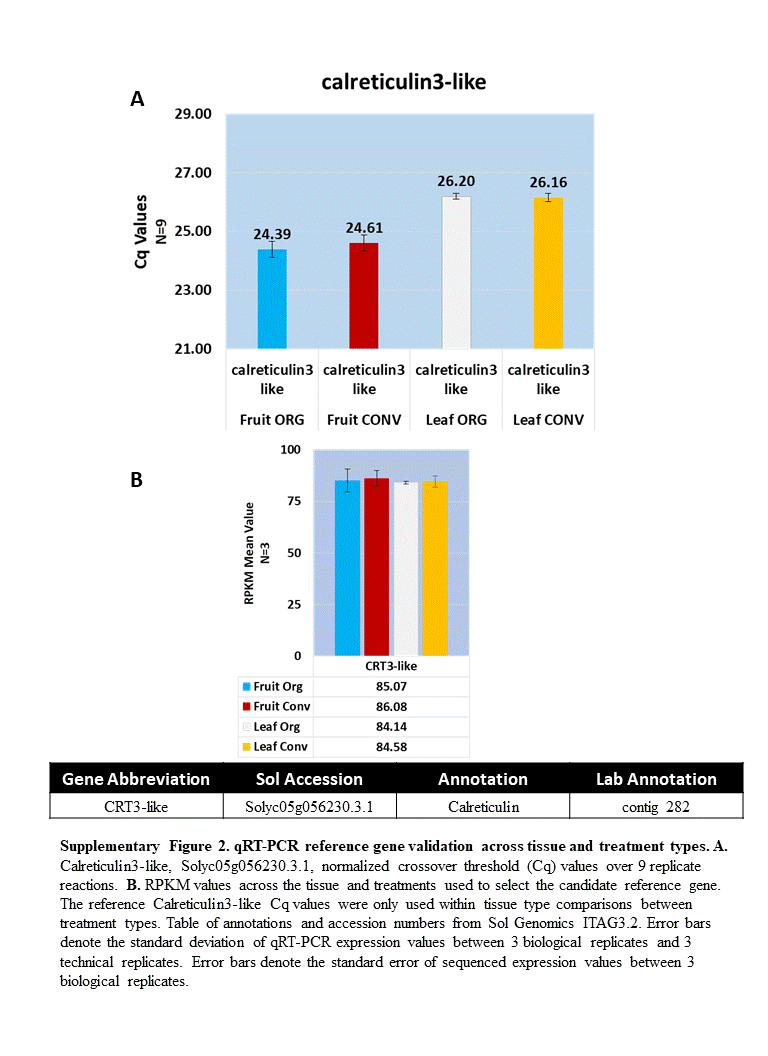

### Supplementary Figure 3A

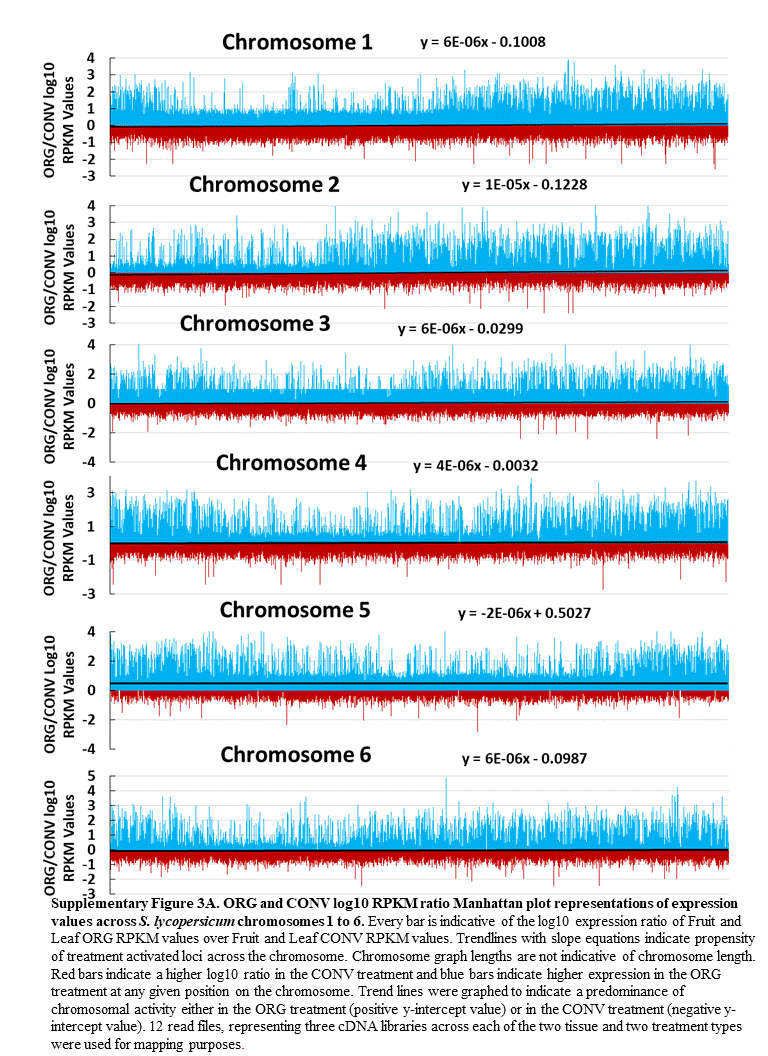

### Supplementary Figure 3B

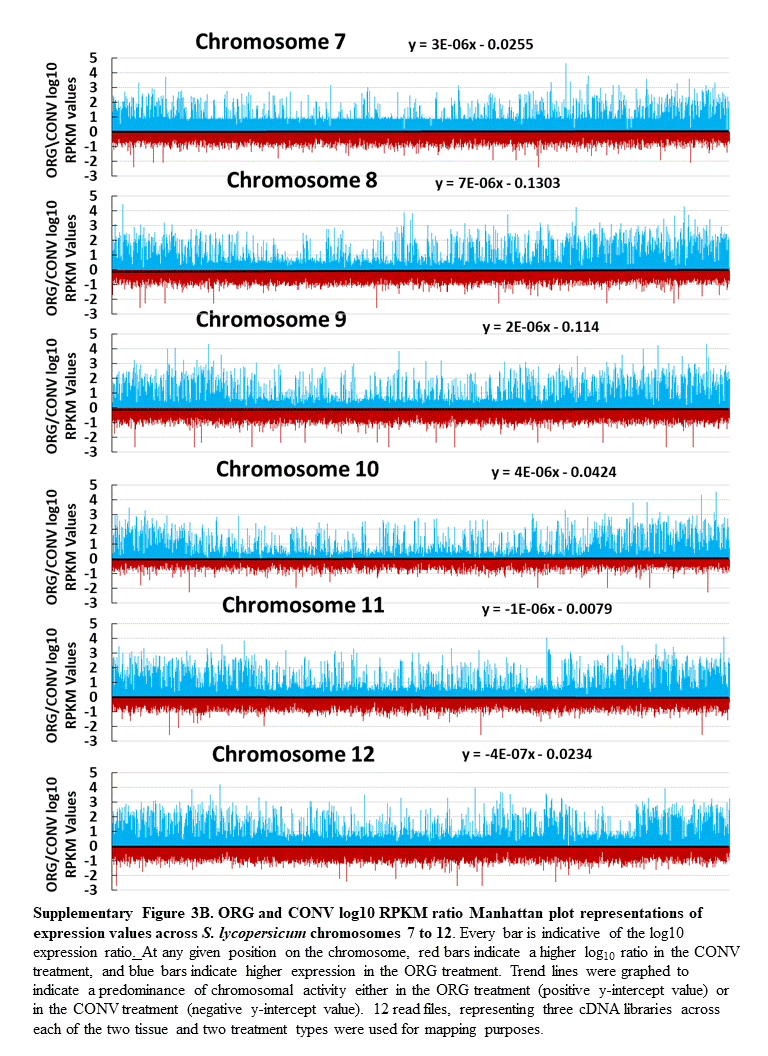

### Supplementary Figure 4

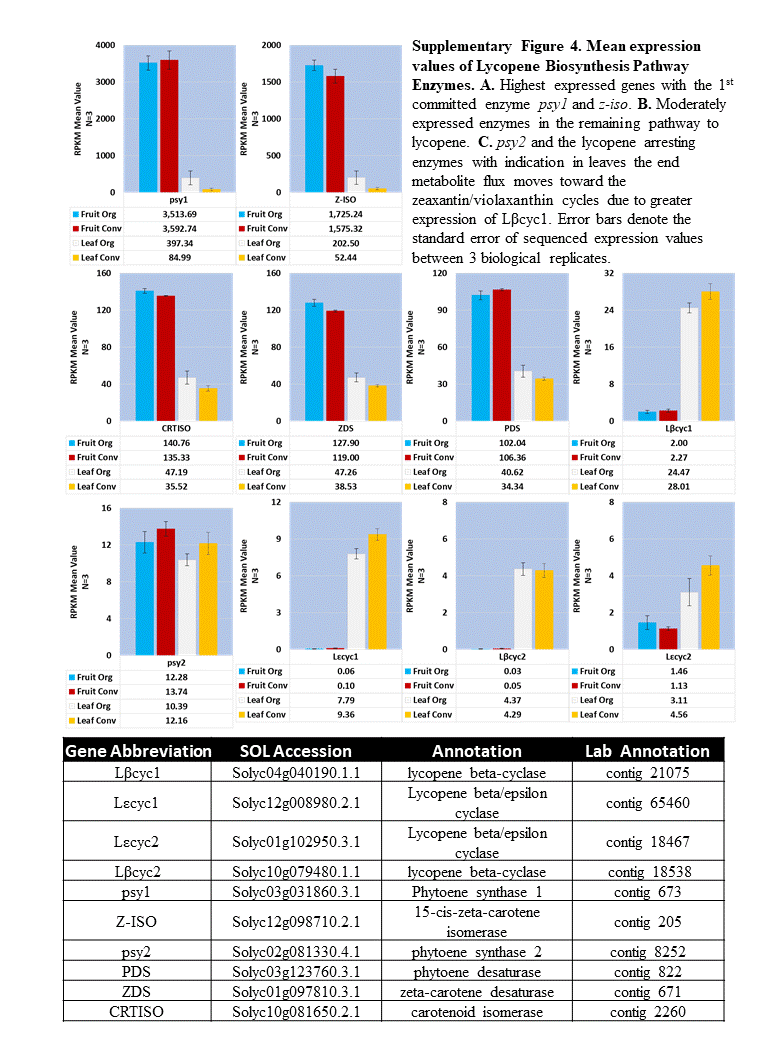

### Supplementary Figure 5

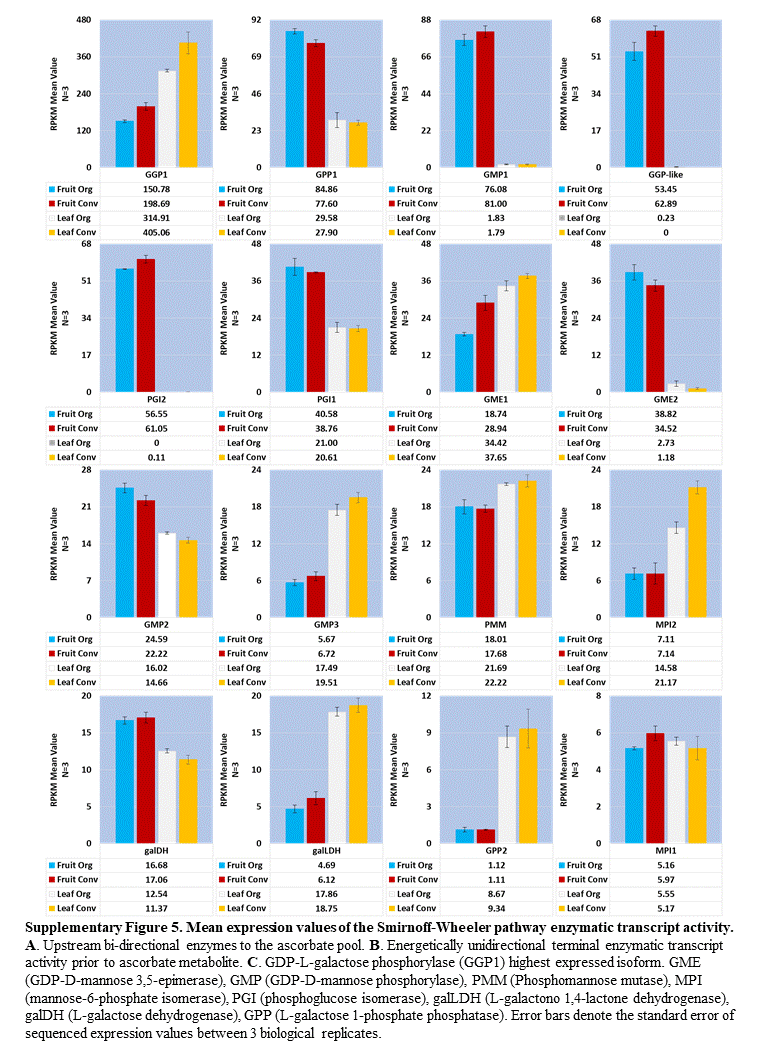

### Supplementary Figure 6A

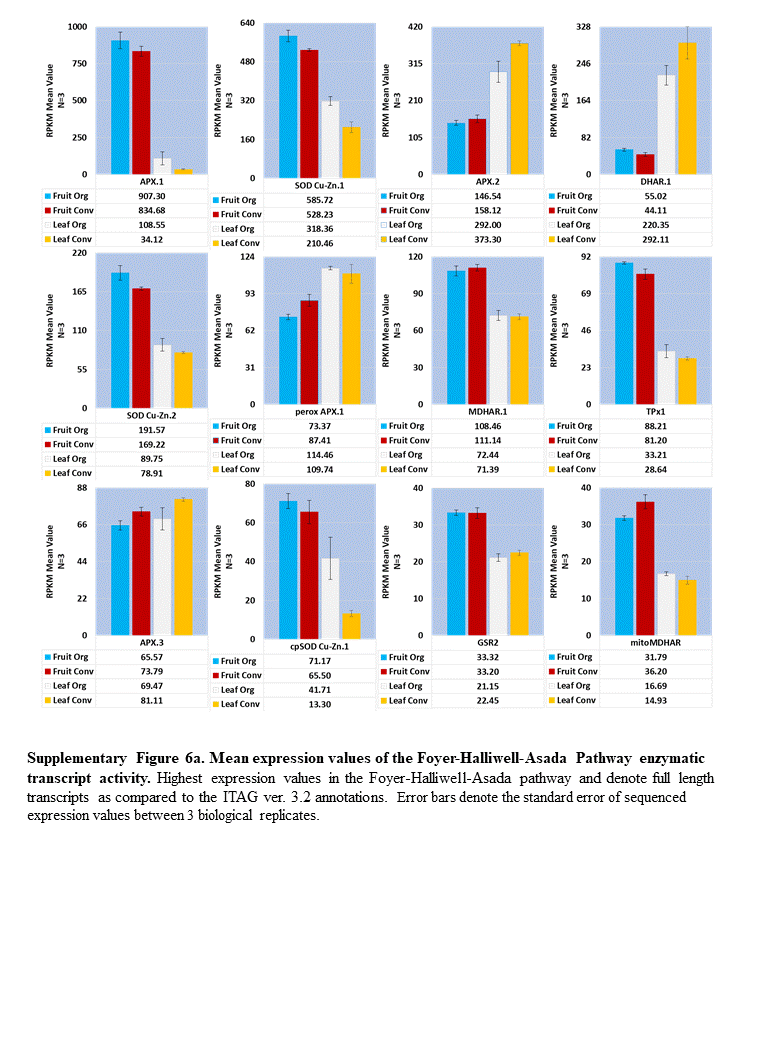

### Supplementary Figure 6B

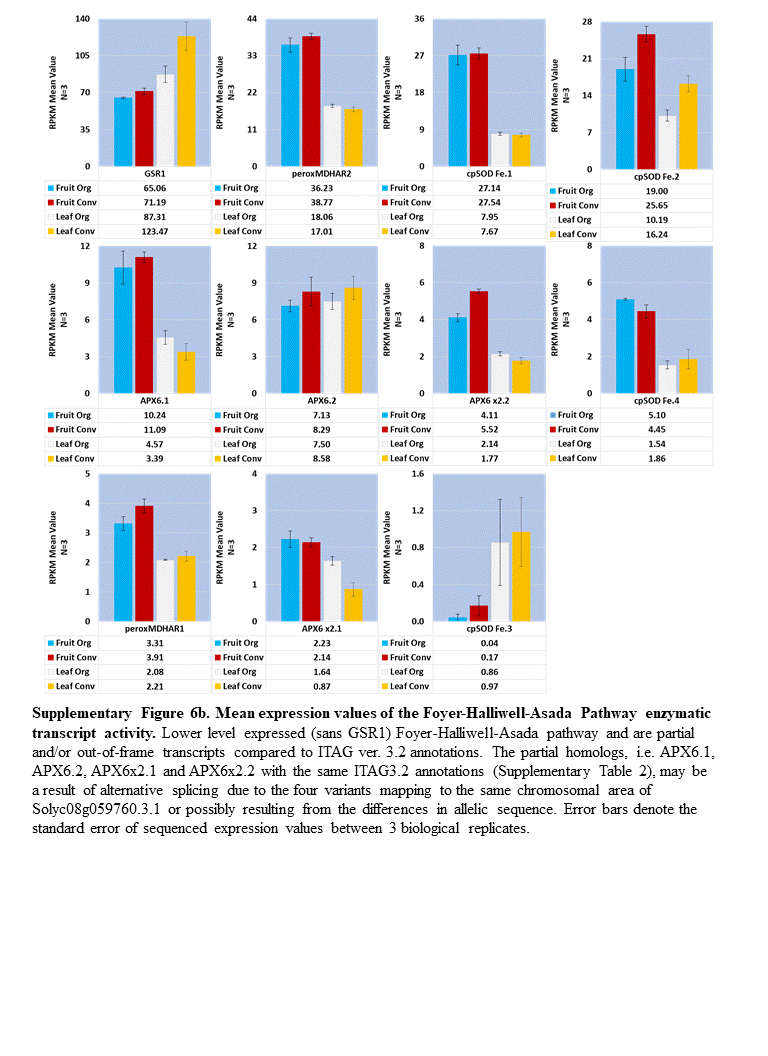

### Supplementary Figure 7

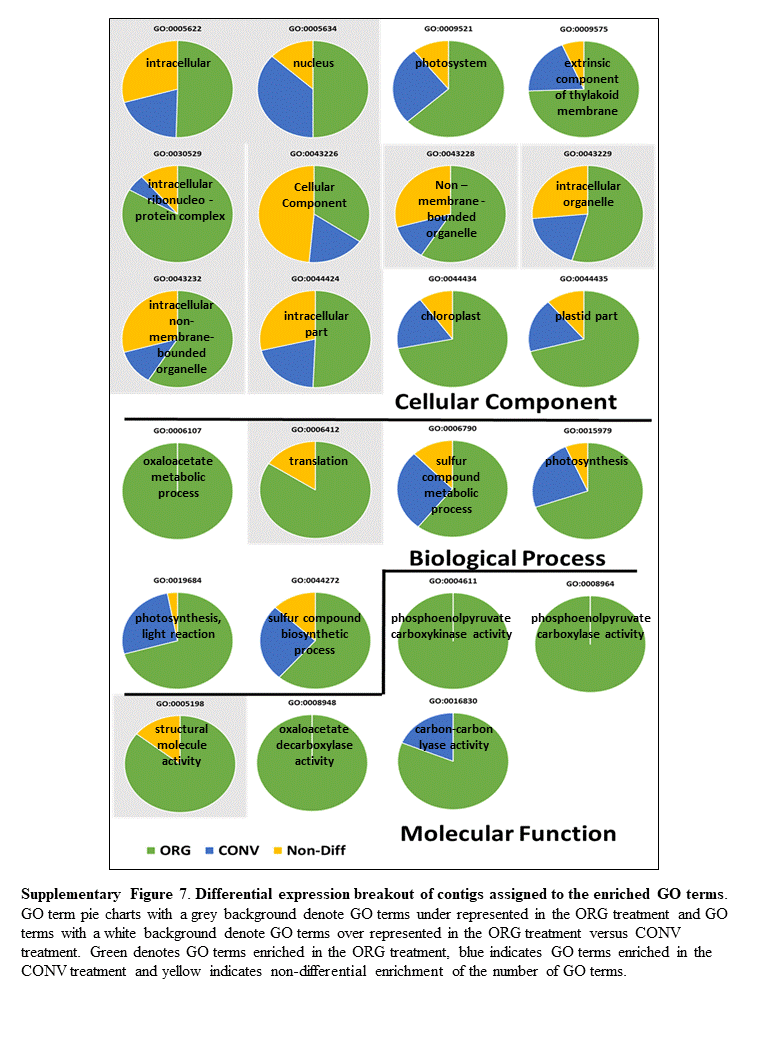

### Supplementary Figure 8

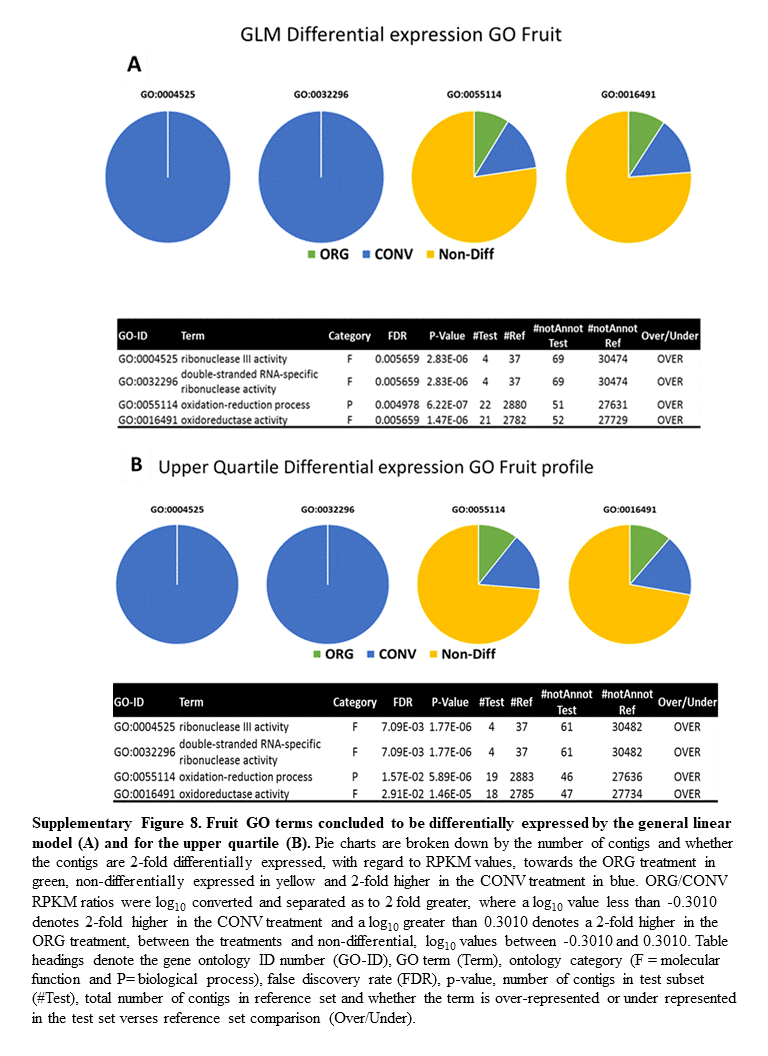

### Supplementary Figure 9A

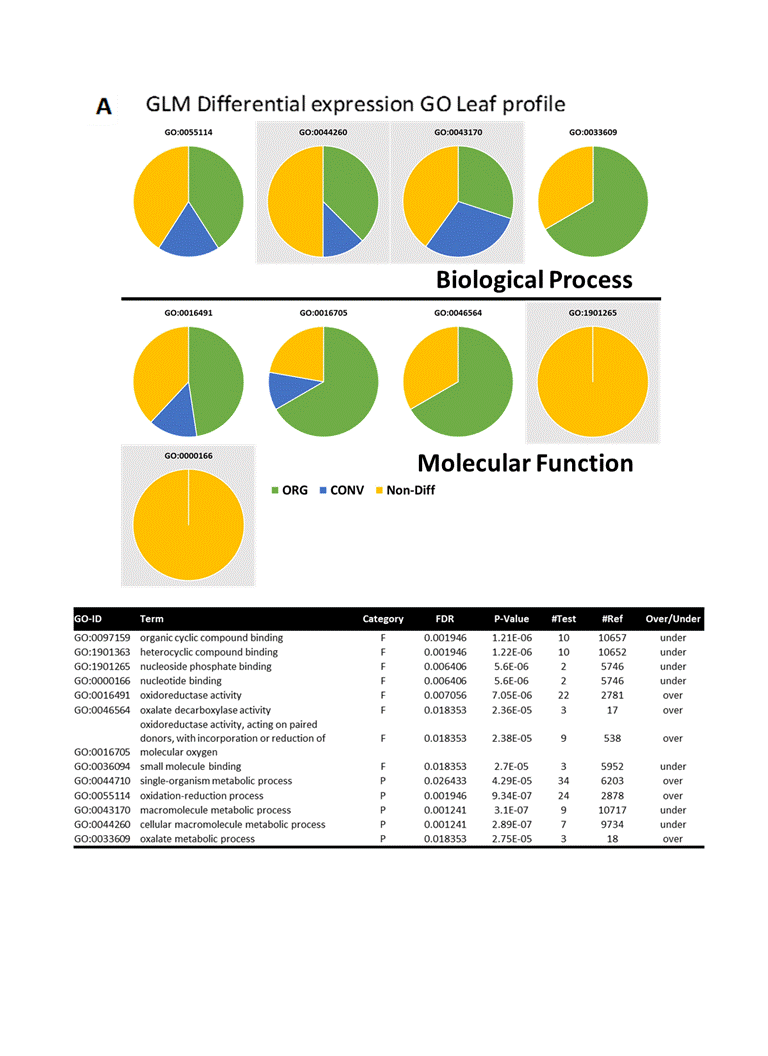

### Supplementary Figure 9B

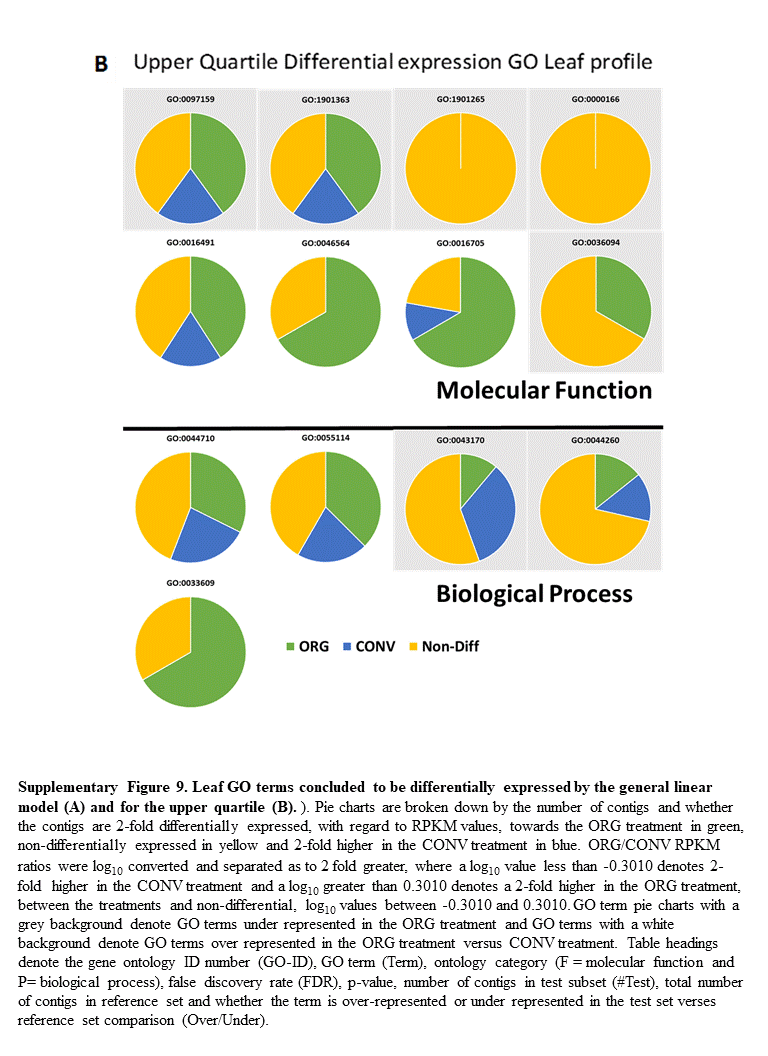

### Supplementary Figure 10

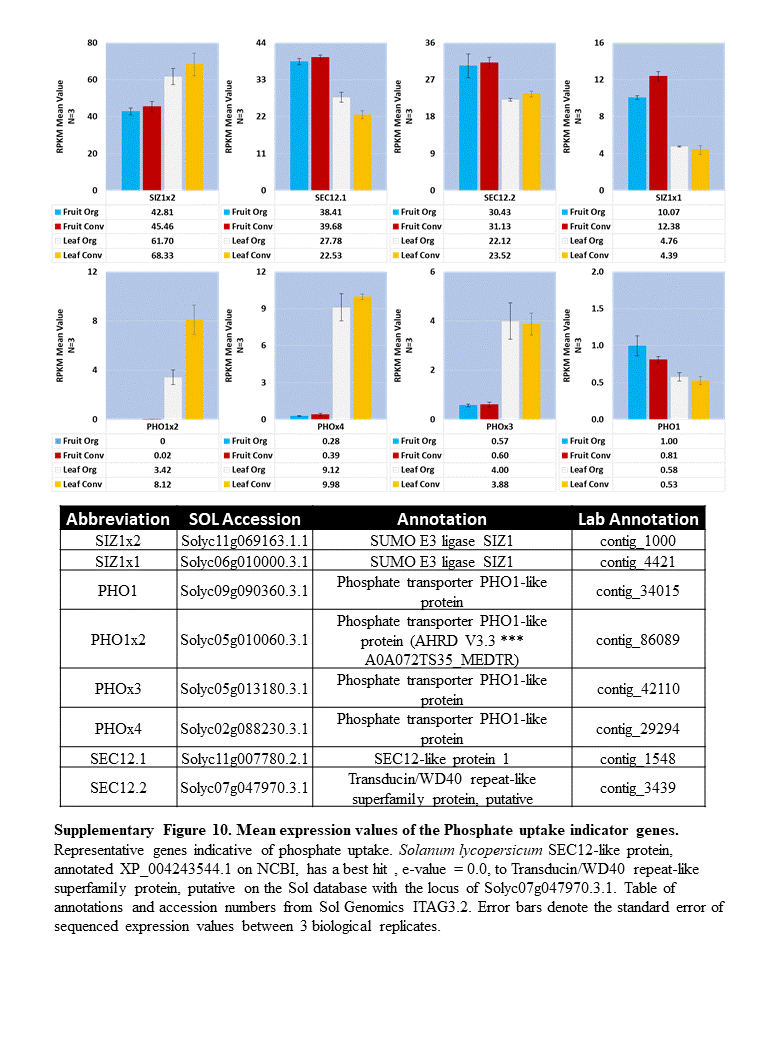

### Supplementary Figure 11

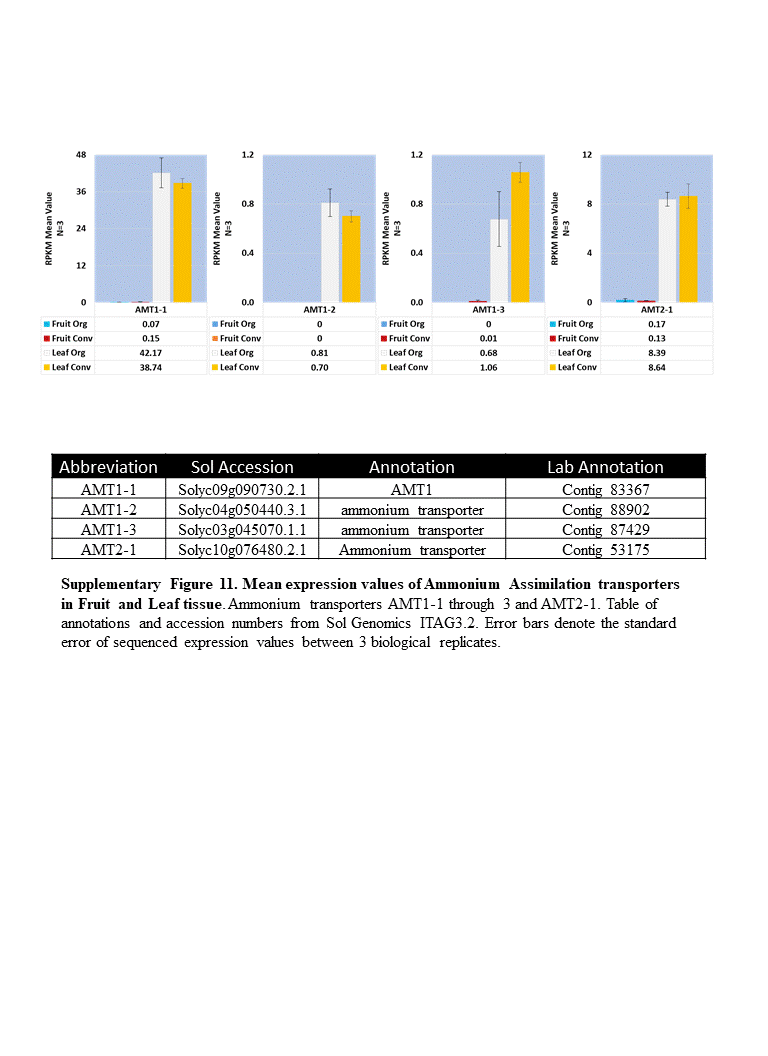

### Supplementary Figure 12

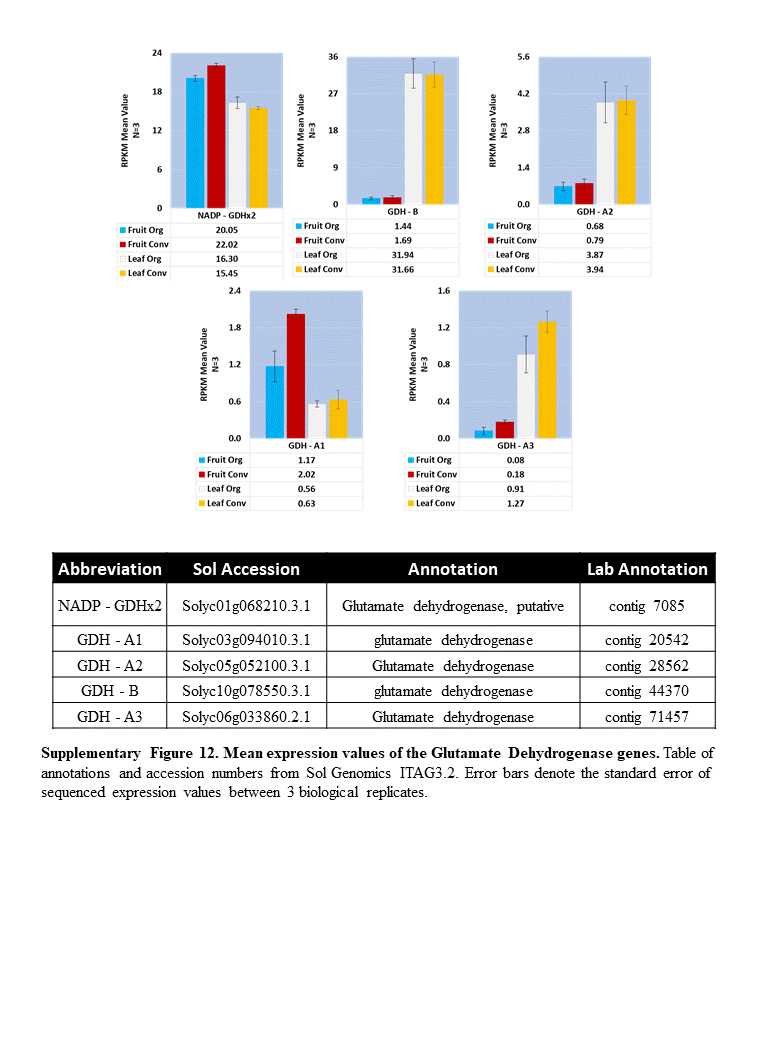

### Supplementary Figure 13A

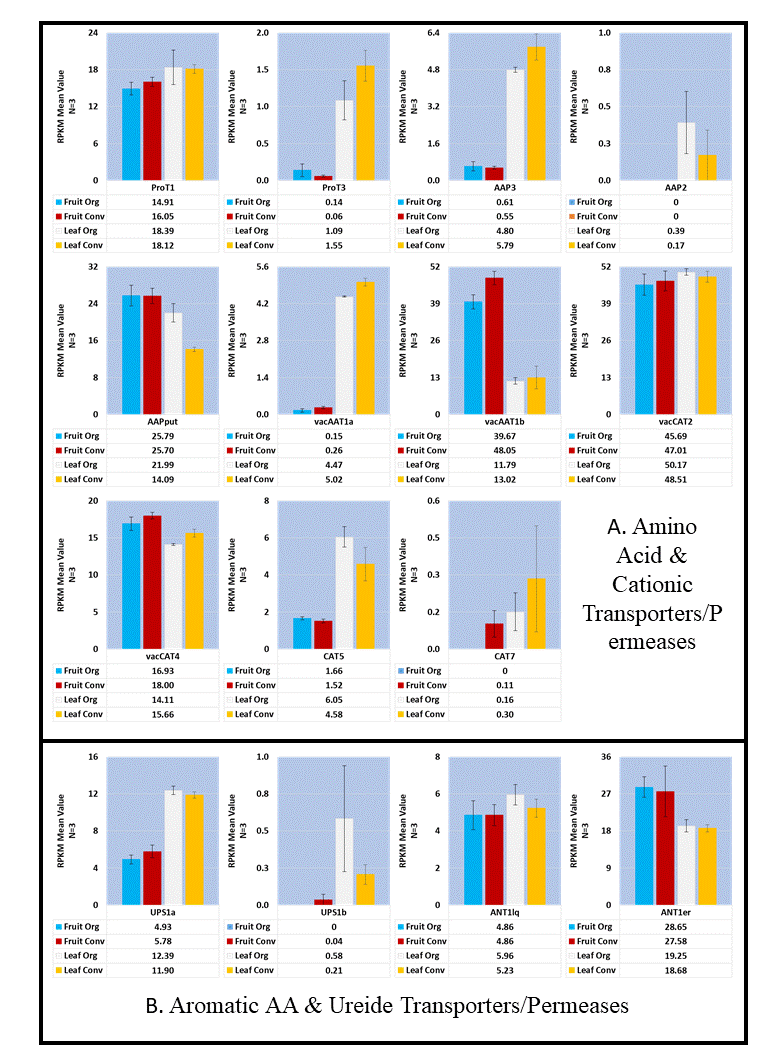

### Supplementary Figure 13B

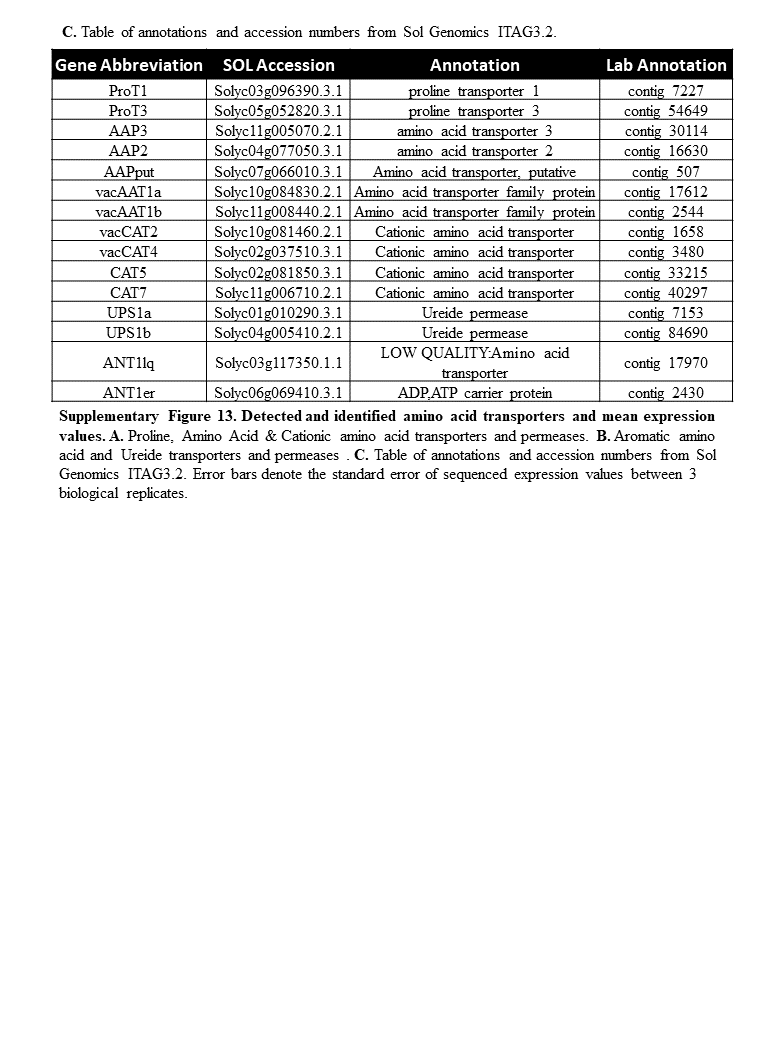

### Supplementary Figure 14

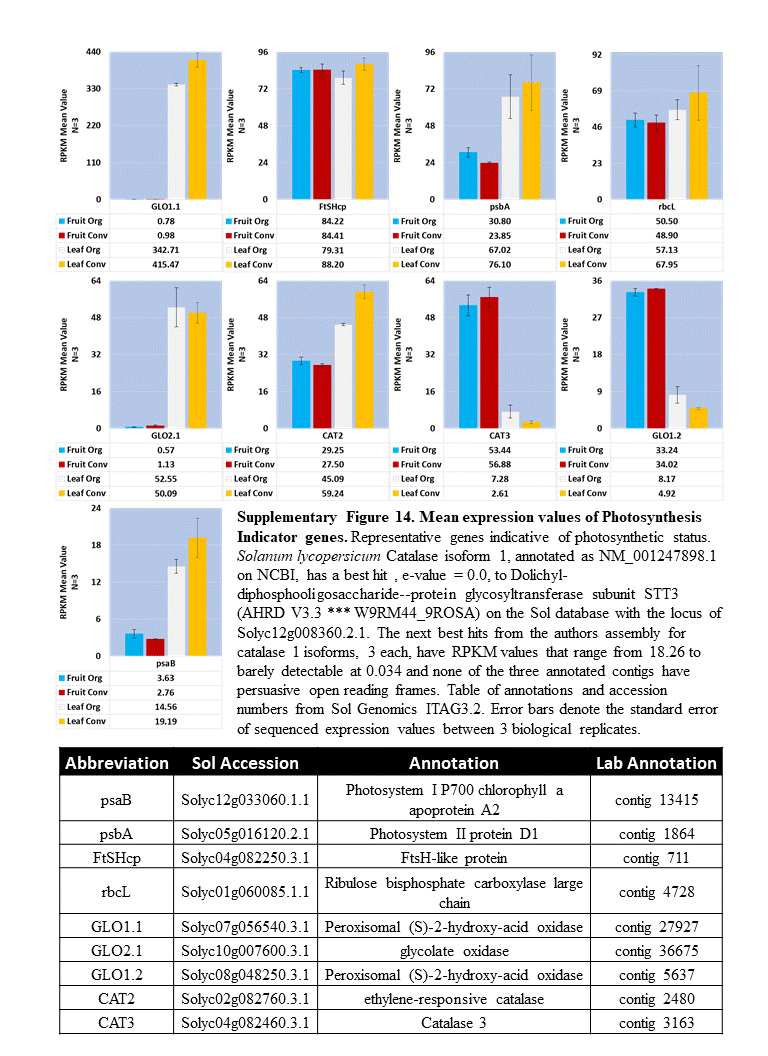

### Supplementary Figure 15A

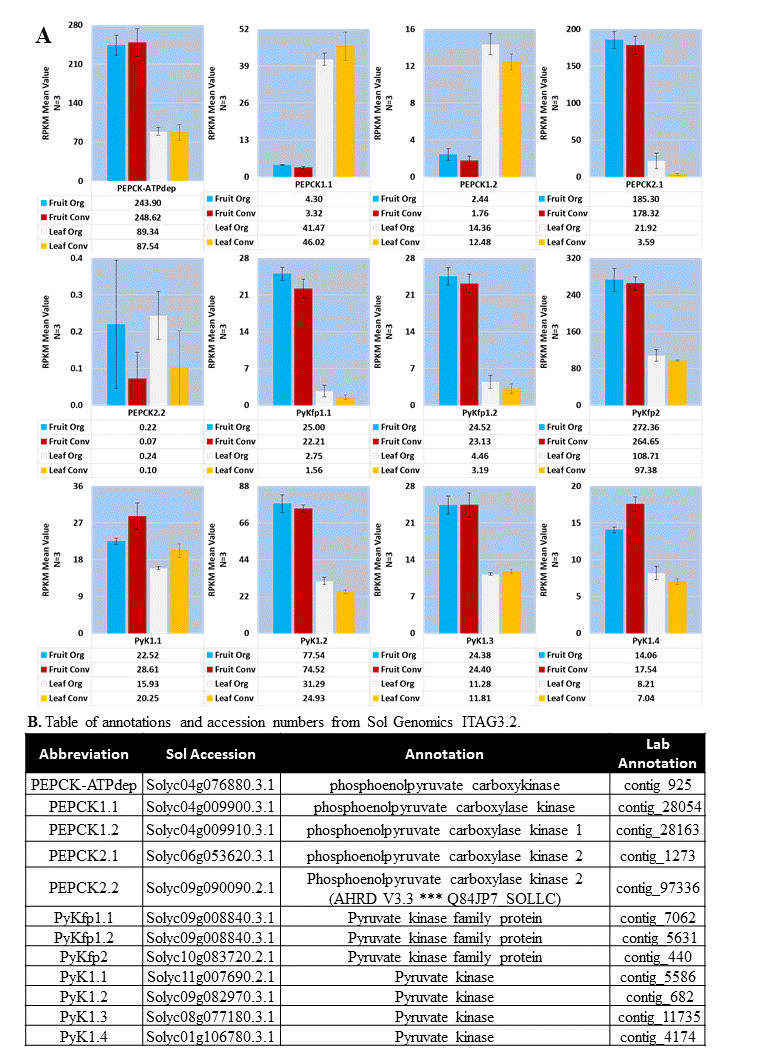

### Supplementary Figure 15B

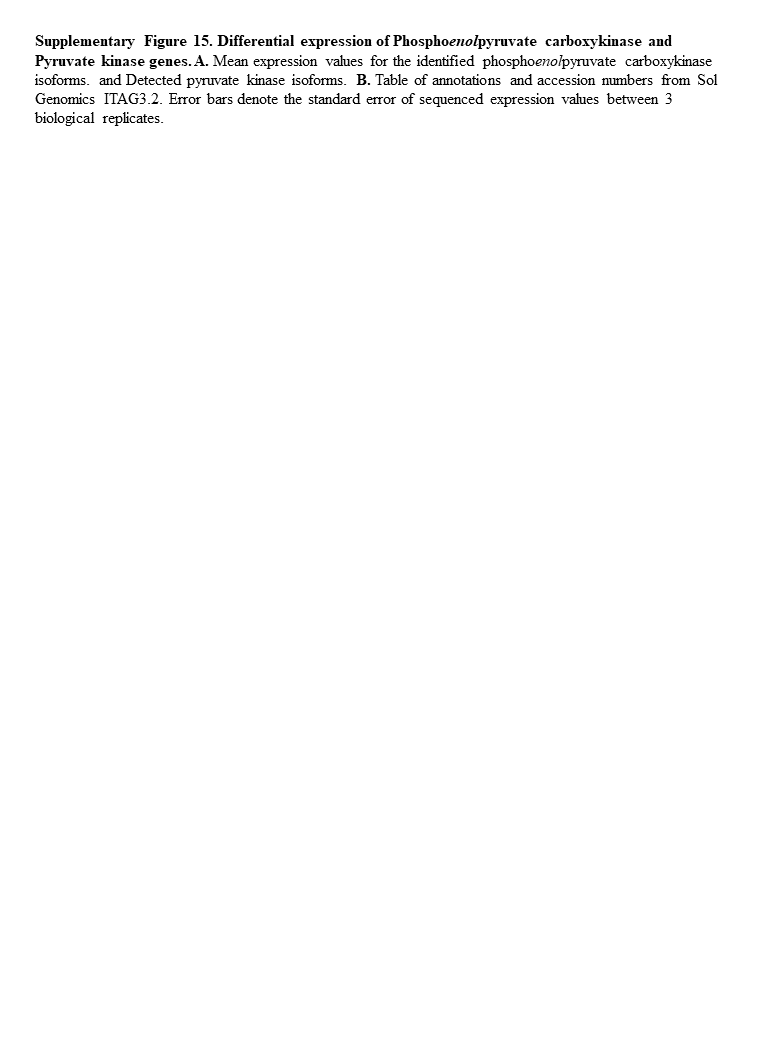

### Supplementary Figure 16A

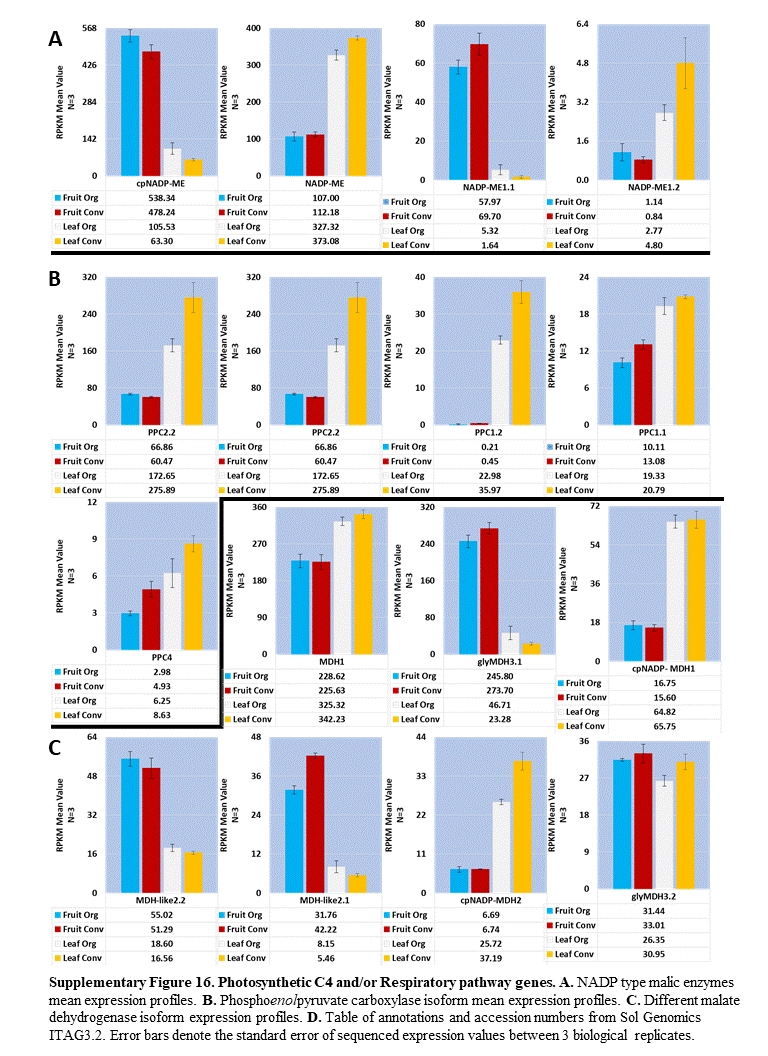

### Supplementary Figure 16B

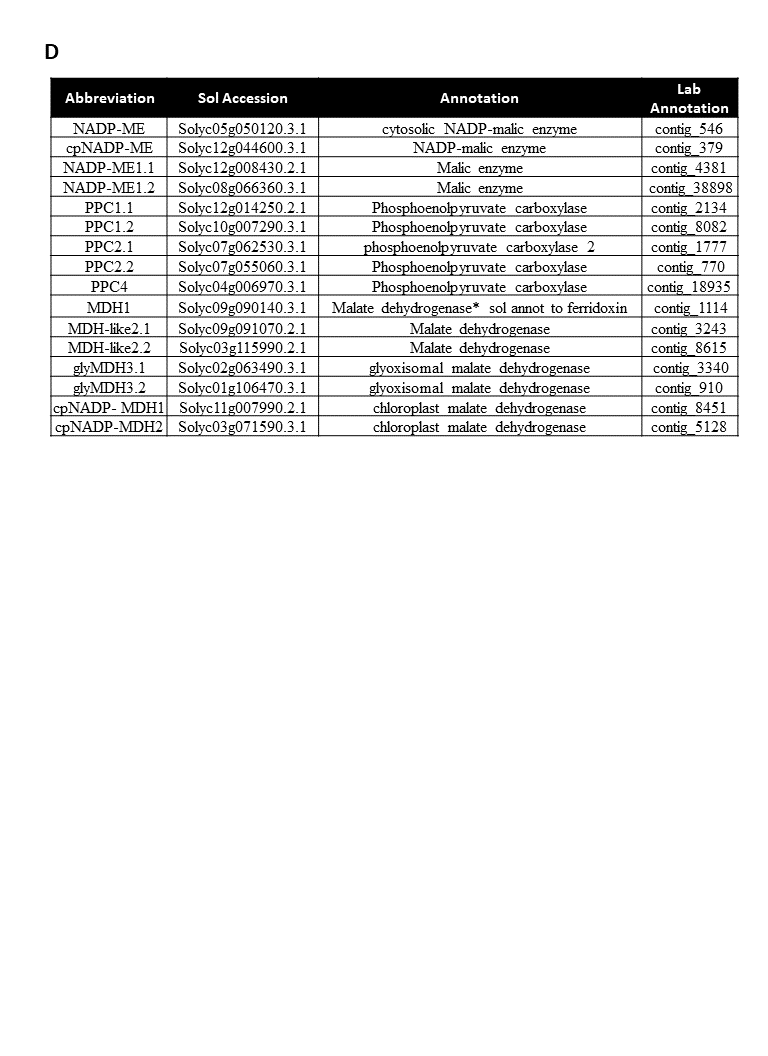

### Supplementary Figure 17A

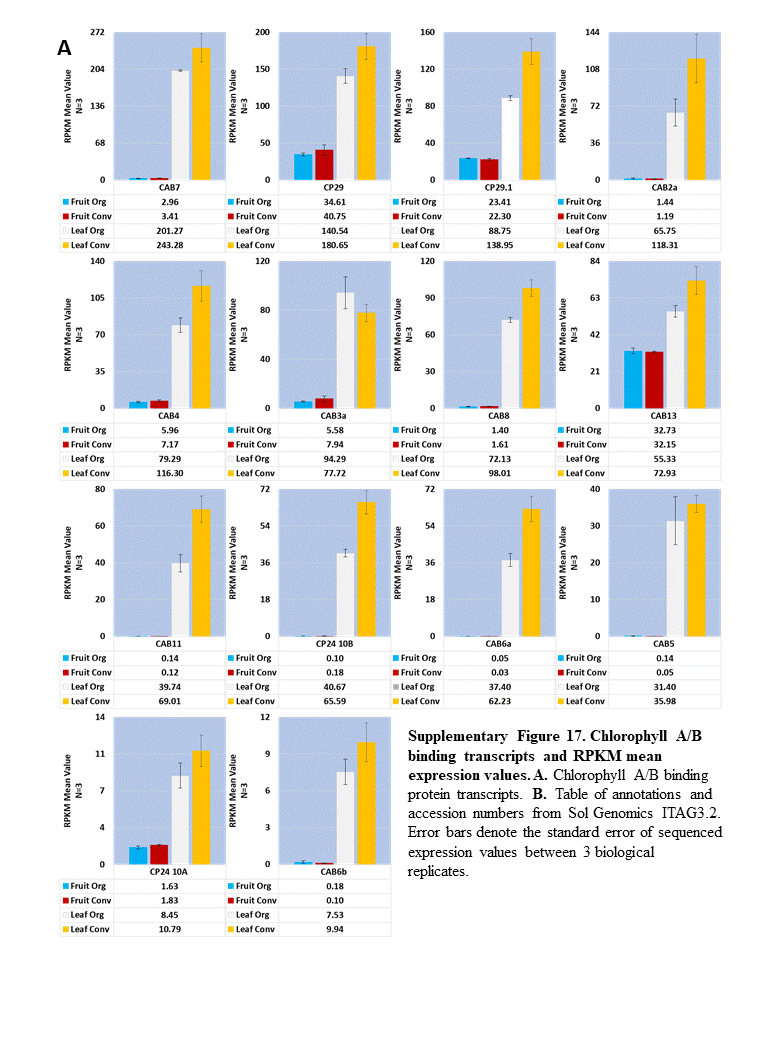

### Supplementary Figure 17B

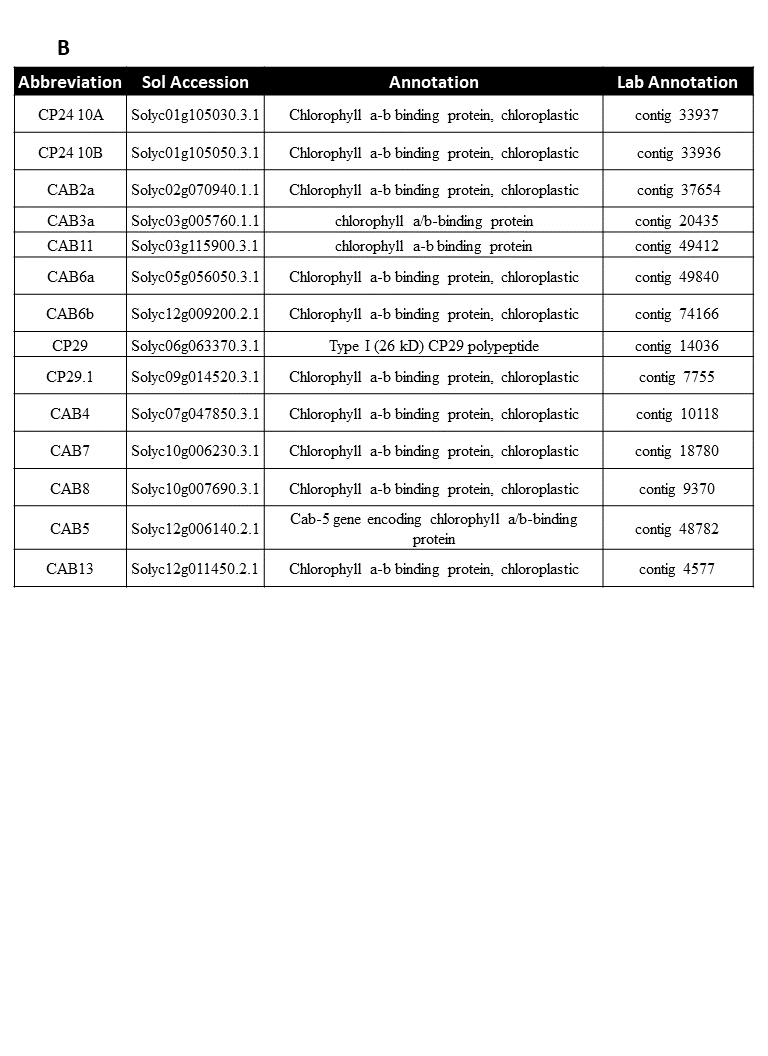

### Supplementary Table 1

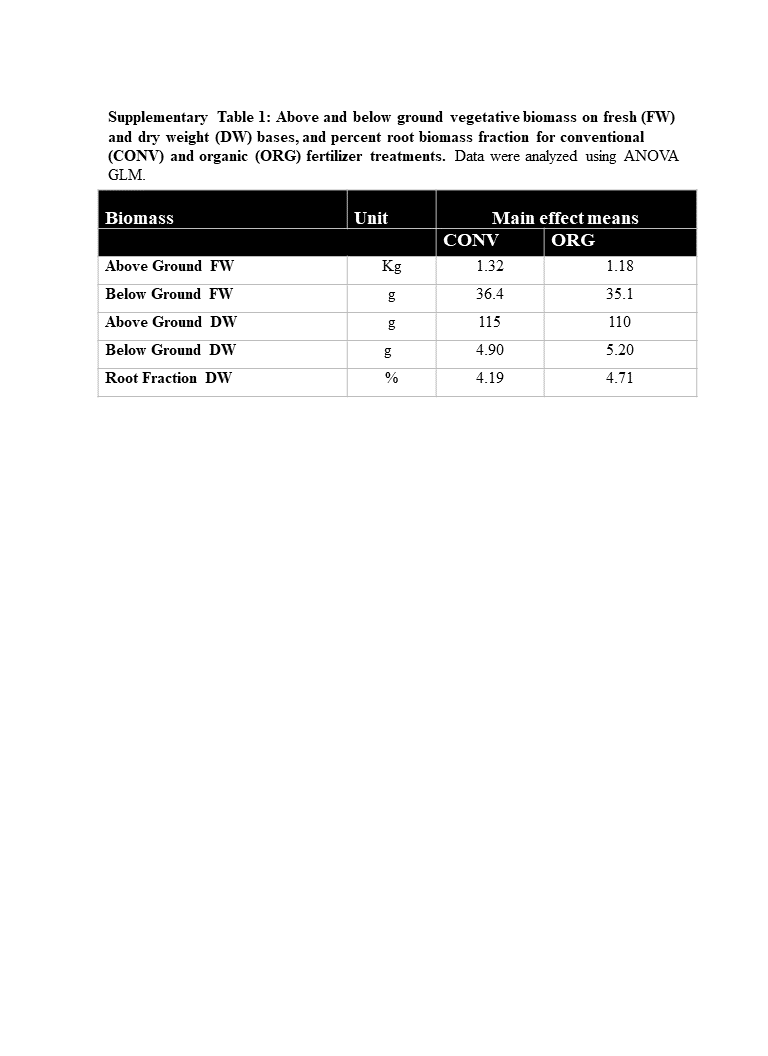

### Supplementary Table 2

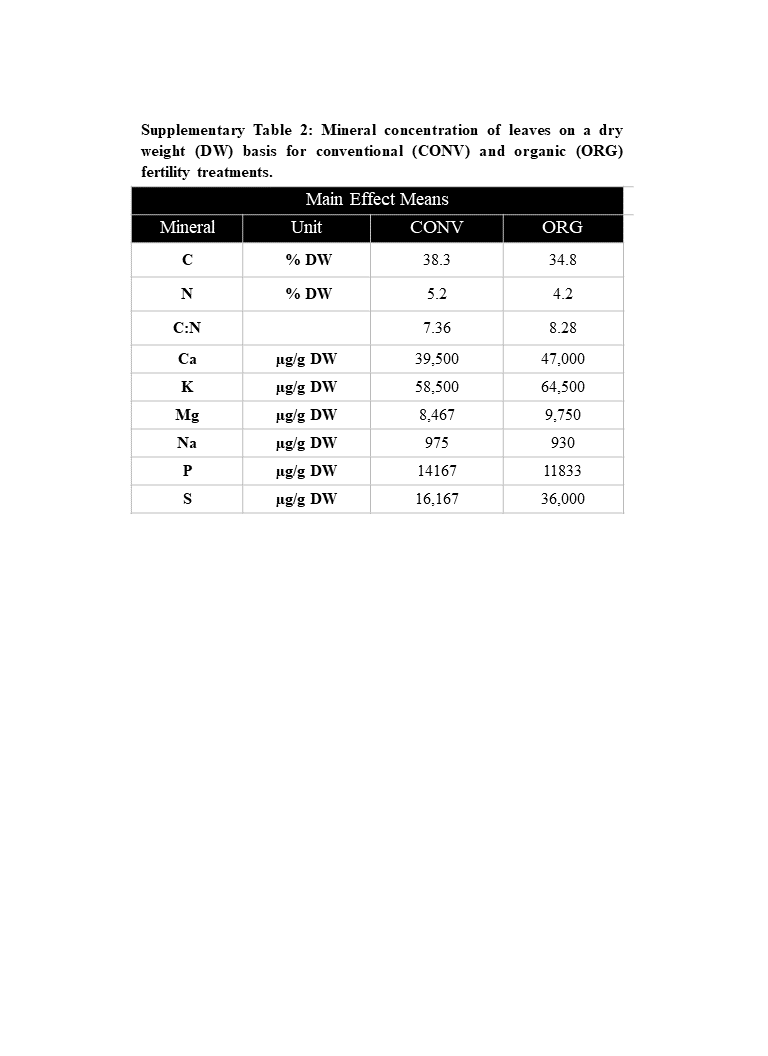

### Supplementary Table 3

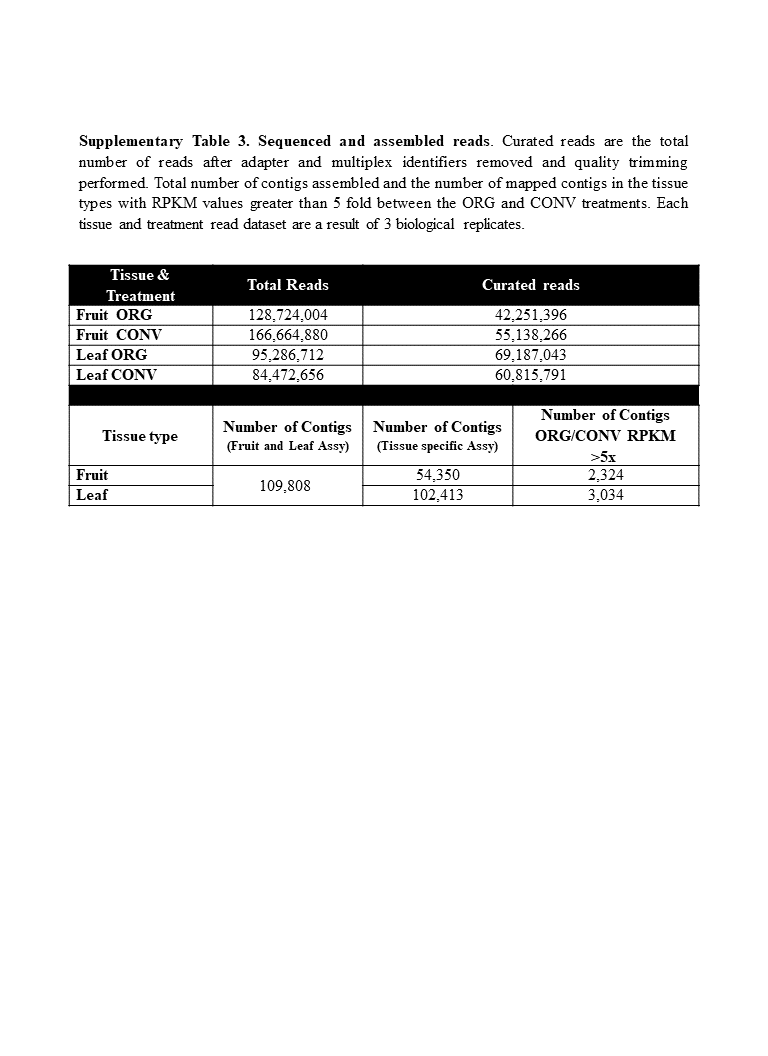

### Supplementary Table 4

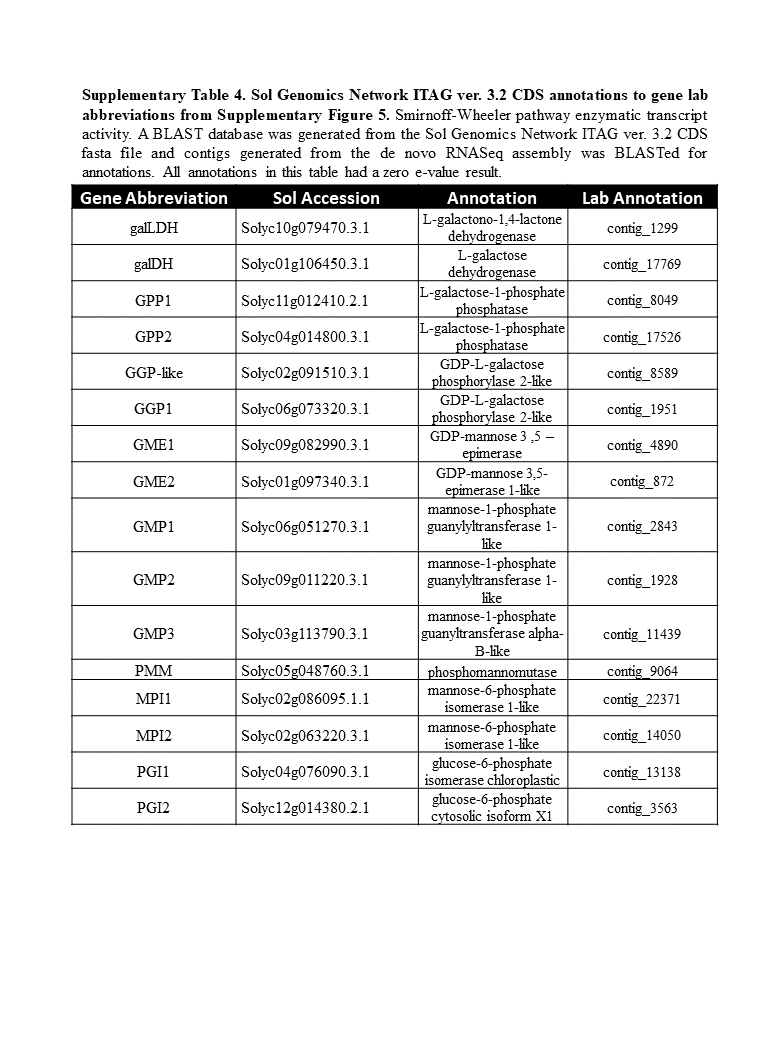

### Supplementary Table 5

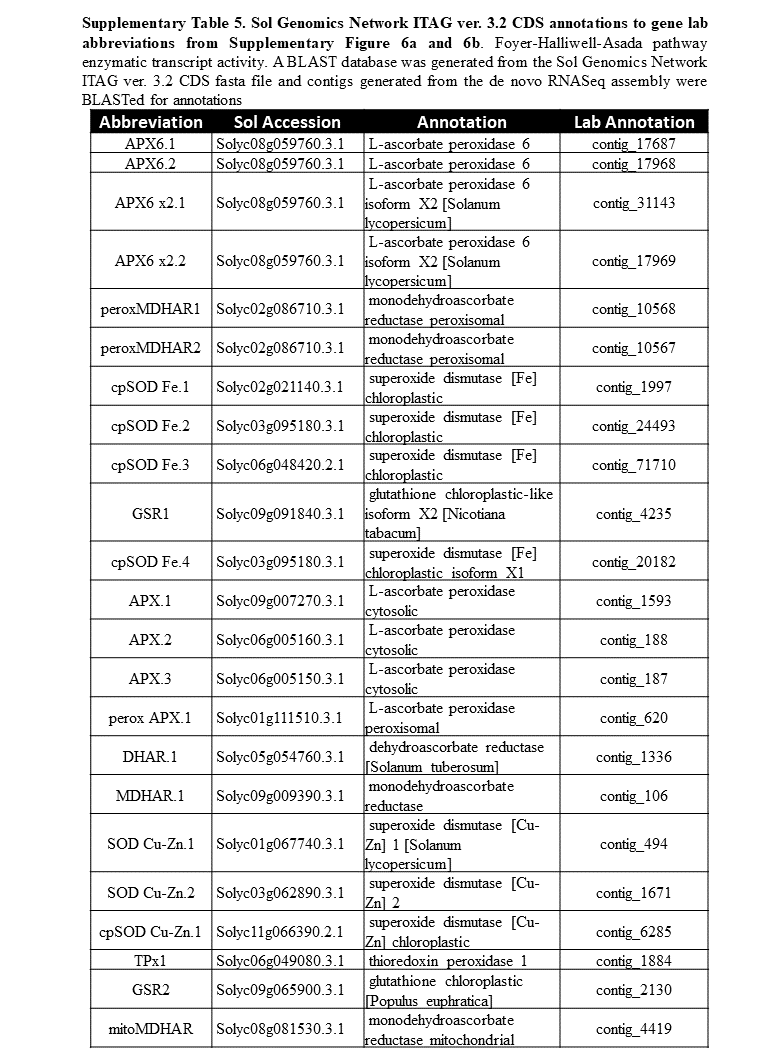
